## Supplementary figures and images for "Single cell RNA-Seq reveals distinct stem cell populations that drive sensory hair cell regeneration in response to loss of Fgf and Notch signaling"

### Supplementary file 3

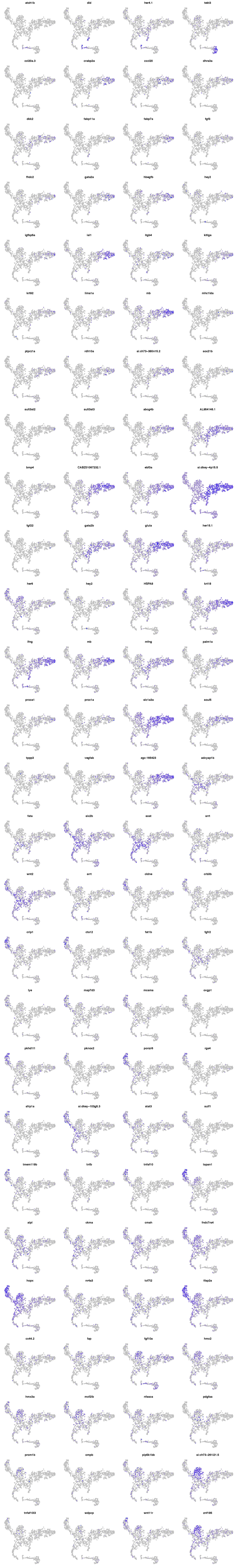

### Supplementary file 5

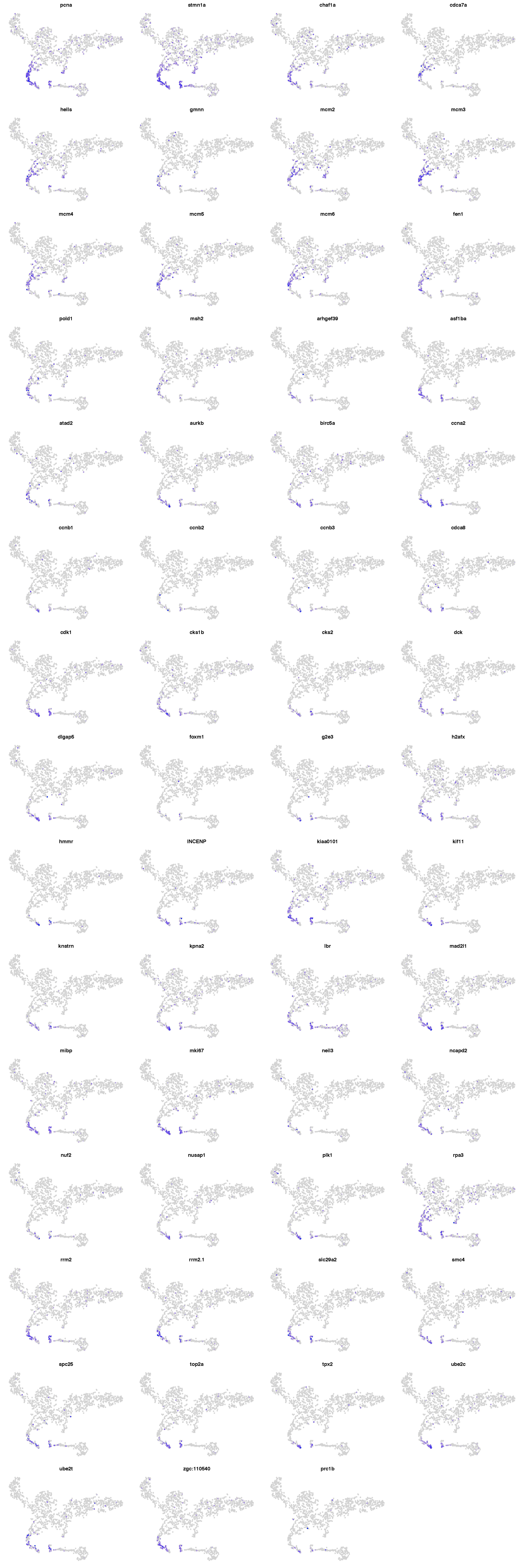

### Supplementary file 10

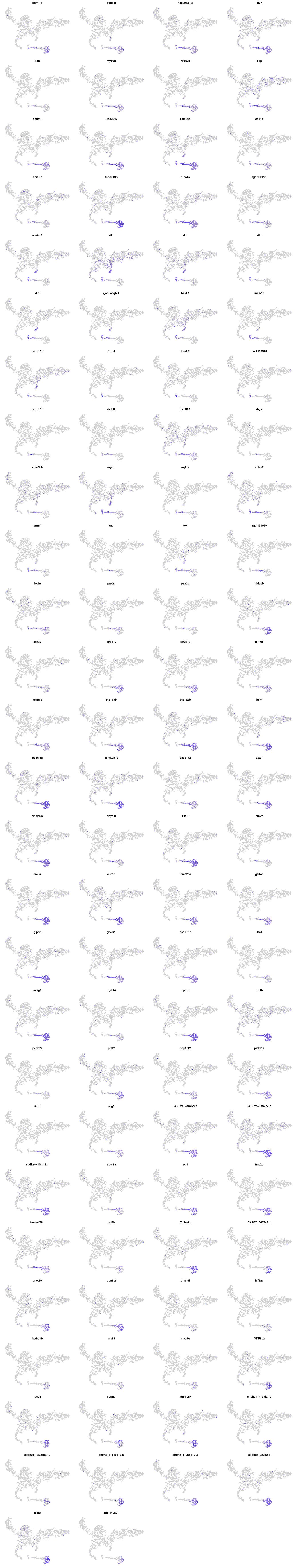
