## Supplementary material for "Single cell RNA-Seq reveals distinct stem cell populations that drive sensory hair cell regeneration in response to loss of Fgf and Notch signaling"

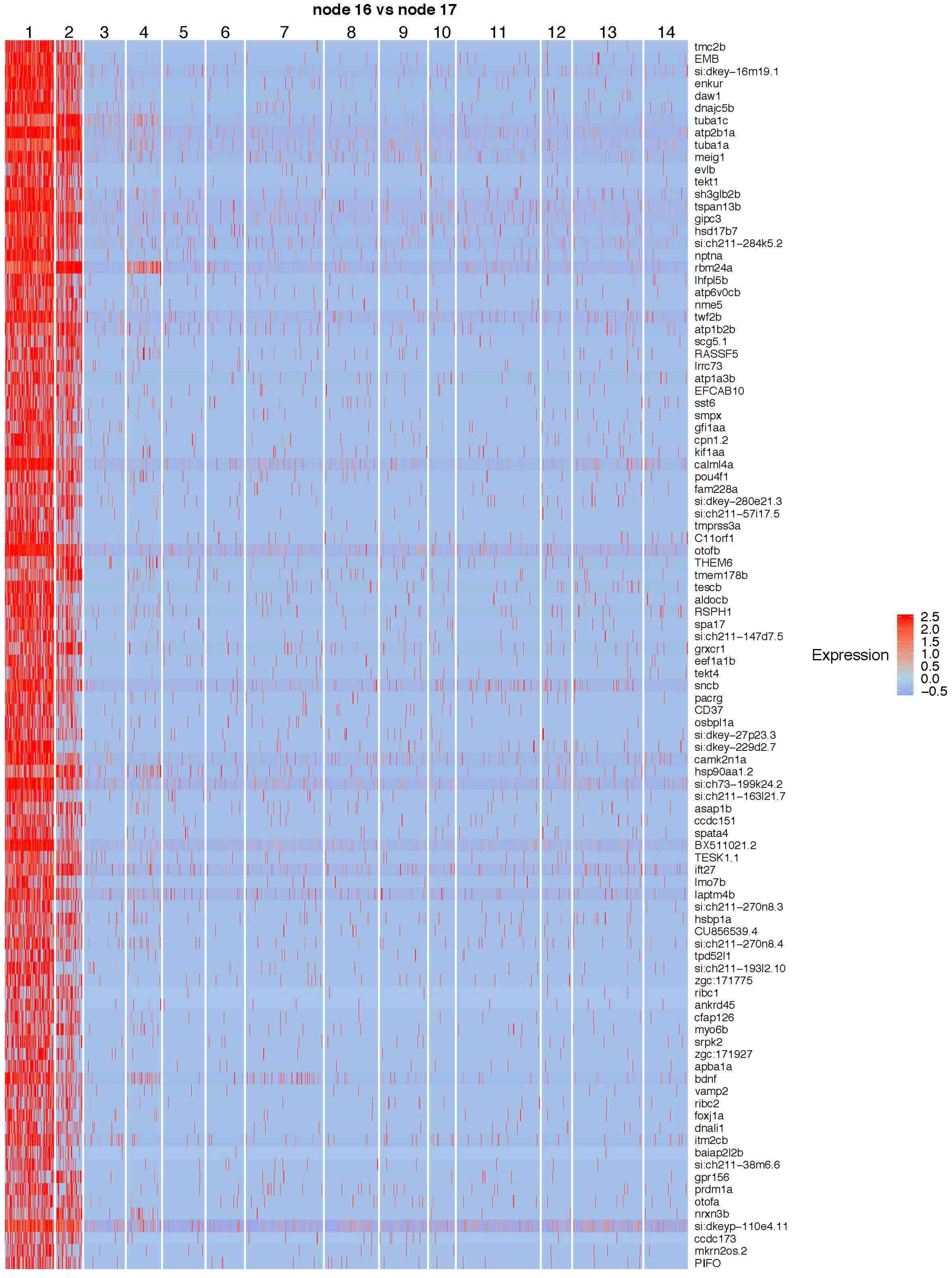
node 16 vs node 17
1
2
3
4
5
6
7
8
9
10
11
12
13
14
tmc2b
EMB
si:dkey-16m19.1
enkur
daw1
dnajc5b
tuba1c
atp2b1a
tuba1a
meig1
evlb
tekt1
sh3glb2b
tspan13b
gipc3
hsd17b7
si:ch211-284k5.2
nptna
rbm24a
lhfpl5b
atp6v0cb
nme5
twf2b
atp1b2b
scg5.1
RASSF5
lrrc73
atp1a3b
EFCAB10
sst6
smpx
gfi1aa
cpn1.2
kif1aa
calml4a
pou4f1
fam228a
si:dkey-280e21.3
si:ch211-57i17.5
tmprss3a
C11orf1
otofb
THEM6
tmem178b
tescb
aldocb
RSPH1
spa17
si:ch211-147d7.5
grxcr1
eef1a1b
tekt4
sncb
pacrg
CD37
osbpl1a
si:dkey-27p23.3
si:dkey-229d2.7
camk2n1a
hsp90aa1.2
si:ch73-199k24.2
si:ch211-163l21.7
asap1b
ccdc151
spata4
BX511021.2
TESK1.1
ift27
lmo7b
laptm4b
si:ch211-270n8.3
hsbp1a
CU856539.4
si:ch211-270n8.4
tpd52l1
si:ch211-193l2.10
zgc:171775
ribc1
ankrd45
cfap126
myo6b
srpk2
zgc:171927
apba1a
bdnf
vamp2
ribc2
foxj1a
dnali1
itm2cb
baiap2l2b
si:ch211-38m6.6
gpr156
prdm1a
otofa
nrxn3b
si:dkeyp-110e4.11
ccdc173
mkrn2os.2
PIFO
Expression
2.5
2.0
1.5
1.0
0.5
0.0
−0.5

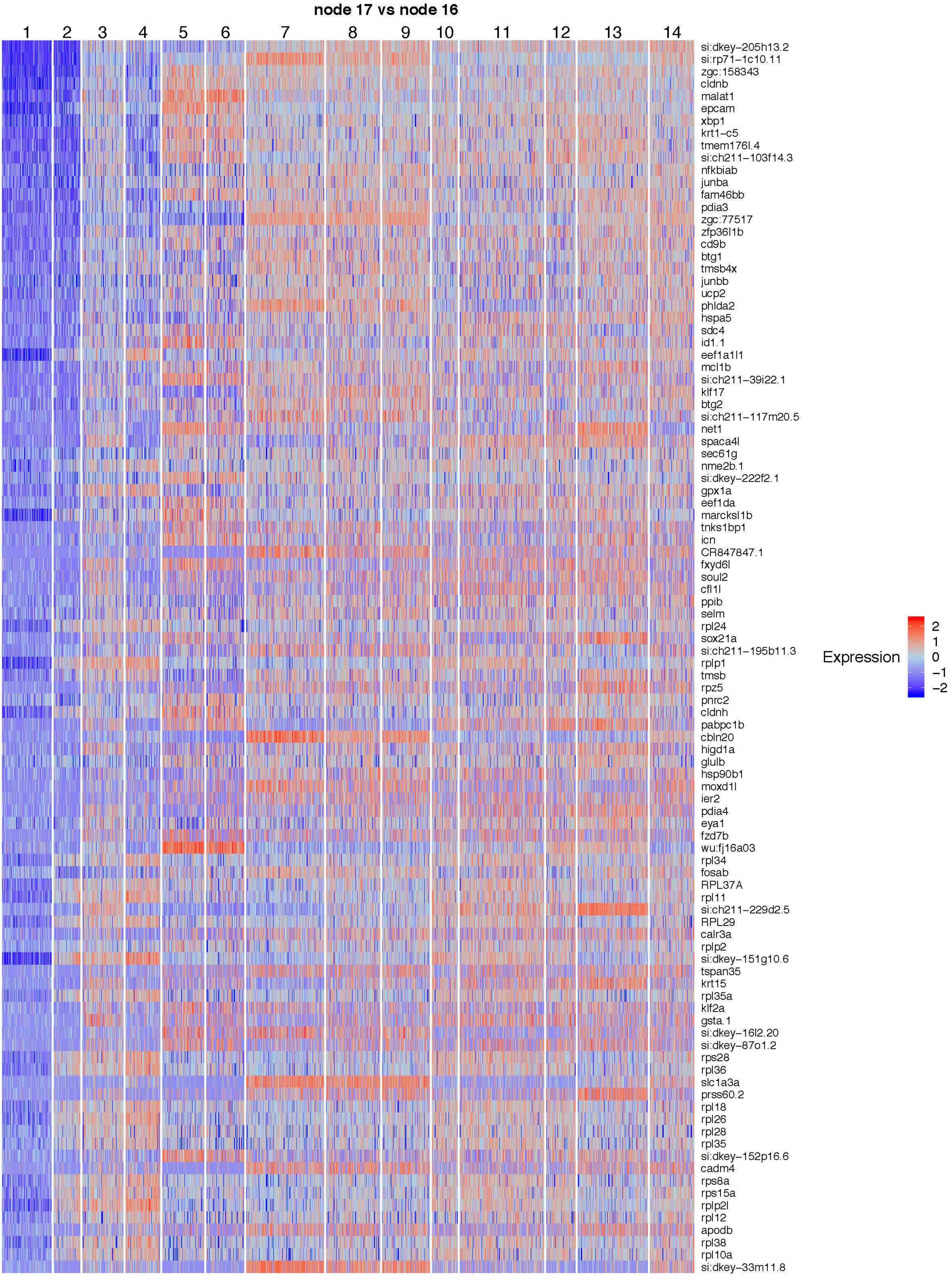
node 17 vs node 16
1
2
3
4
5
6
7
8
9
10
11
12
13
14
si:dkey−205h13.2
si:rp71−1c10.11
zgc:158343
cldnb
malat1
epcam
xbp1
krt1−c5
tmem176l.4
si:ch211−103f14.3
nfkbiab
junba
fam46bb
pdia3
zgc:77517
zfp36l1b
cd9b
btg1
tmsb4x
junbb
ucp2
phlda2
hspa5
sdc4
id1.1
eef1a1l1
mcl1b
si:ch211−39i22.1
klf17
btg2
si:ch211−117m20.5
net1
spaca4l
sec61g
nme2b.1
si:dkey−222f2.1
gpx1a
eef1da
marcksl1b
tnks1bp1
icn
CR847847.1
fxyd6l
soul2
cfl1l
ppib
selm
rpl24
sox21a
si:ch211−195b11.3
rplp1
tmsb
rpz5
pnrc2
cldnh
pabpc1b
cbln20
higd1a
glulb
hsp90b1
moxd1l
ier2
pdia4
eya1
fzd7b
wu:fj16a03
rpl34
fosab
RPL37A
rpl11
si:ch211−229d2.5
RPL29
calr3a
rplp2
si:dkey−151g10.6
tspan35
krt15
rpl35a
klf2a
gsta.1
si:dkey−16l2.20
si:dkey−87o1.2
rps28
rpl36
slc1a3a
prss60.2
rpl18
rpl26
rpl28
rpl35
si:dkey−152p16.6
cadm4
rps8a
rps15a
rplp2l
rpl12
apodb
rpl38
rpl10a
si:dkey−33m11.8
Expression
2
1
0
−1
−2

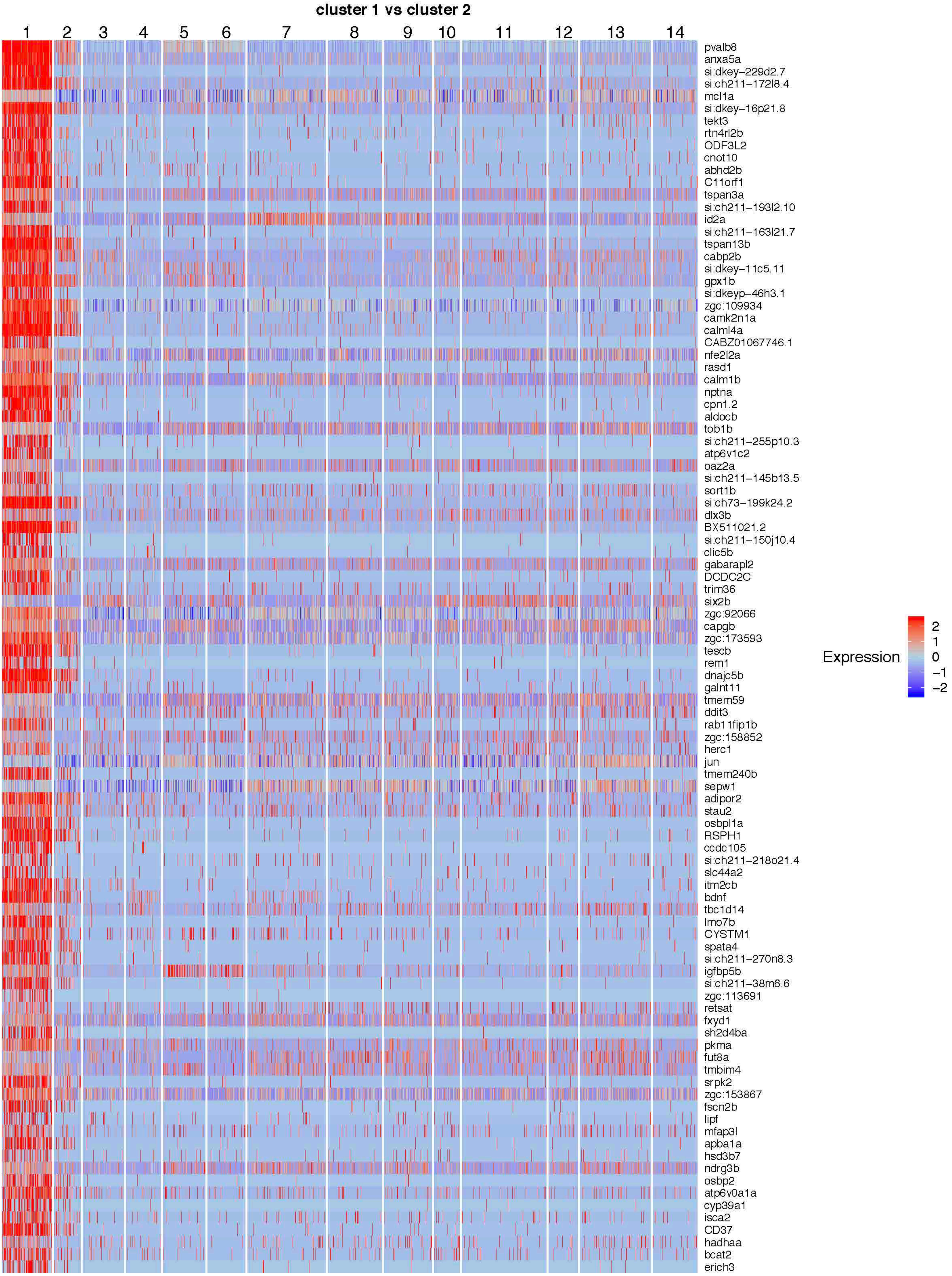
cluster 1 vs cluster 2
1
2
3
4
5
6
7
8
9
10
11
12
13
14
pvalb8
anxa5a
si:dkey-229d2.7
si:ch211-172l8.4
mcl1a
si:dkey-16p21.8
tekt3
rtn4rl2b
ODF3L2
cnot10
abhd2b
C11orf1
tspan3a
si:ch211-193l2.10
id2a
si:ch211-163l21.7
tspan13b
cabp2b
si:dkey-11c5.11
gpx1b
si:dkeyp-46h3.1
zgc:109934
camk2n1a
calml4a
CABZ01067746.1
nfe2l2a
rasd1
calm1b
nptna
cpn1.2
aldocb
tob1b
si:ch211-255p10.3
atp6v1c2
oaz2a
si:ch211-145b13.5
sort1b
si:ch73-199k24.2
dlx3b
BX511021.2
si:ch211-150j10.4
clic5b
gabarapl2
DCDC2C
trim36
six2b
zgc:92066
capgb
zgc:173593
tescb
rem1
dnajc5b
galnt11
tmem59
ddit3
rab11fip1b
zgc:158852
herc1
jun
tmem240b
sepw1
adipor2
stau2
osbpl1a
RSPH1
ccdc105
si:ch211-218o21.4
slc44a2
itm2cb
bdnf
tbc1d14
lmo7b
CYSTM1
spata4
si:ch211-270n8.3
igfbp5b
si:ch211-38m6.6
zgc:113691
retsat
fxyd1
sh2d4ba
pkma
fut8a
tmbim4
srpk2
zgc:153867
fscn2b
lipf
mfap3l
apba1a
hsd3b7
ndrg3b
osbp2
atp6v0a1a
cyp39a1
isca2
CD37
hadhaa
bcat2
erich3
Expression
2
1
0
−1
−2

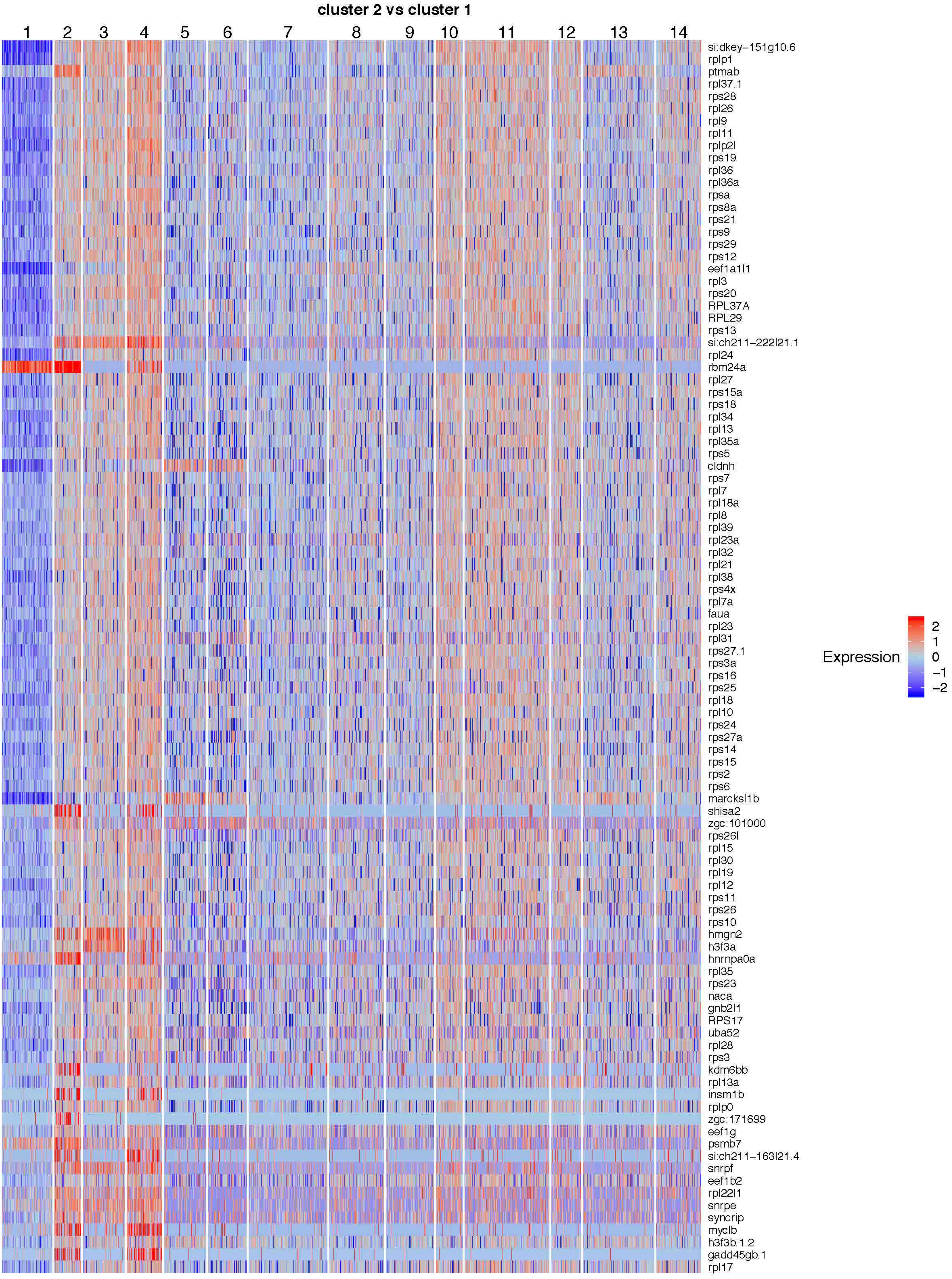
cluster 2 vs cluster 1
1
2
3
4
5
6
7
8
9
10
11
12
13
14
si:dkey−151g10.6
rplp1
ptmab
rpl37.1
rps28
rpl26
rpl9
rpl11
rplp2l
rps19
rpl36
rpl36a
rpsa
rps8a
rps21
rps9
rps29
rps12
eef1a1l1
rpl3
rps20
RPL37A
RPL29
rps13
si:ch211−222l21.1
rpl24
rbm24a
rpl27
rps15a
rps18
rpl34
rpl13
rpl35a
rps5
cldnh
rps7
rpl7
rpl18a
rpl8
rpl39
rpl23a
rpl32
rpl21
rpl38
rps4x
rpl7a
faua
rpl23
rpl31
rps27.1
rps3a
rps16
rps25
rpl18
rpl10
rps24
rps27a
rps14
rps15
rps2
rps6
marcksl1b
shisa2
zgc:101000
rps26l
rpl15
rpl30
rpl19
rpl12
rps11
rps26
rps10
hmgn2
h3f3a
hnrnpa0a
rpl35
rps23
naca
gnb2l1
RPS17
uba52
rpl28
rps3
kdm6bb
rpl13a
insm1b
rplp0
zgc:171699
eef1g
psmb7
si:ch211−163l21.4
snrpf
eef1b2
rpl22l1
snrpe
syncrip
myclb
h3f3b.1.2
gadd45gb.1
rpl17
Expression
2
1
0
−1
−2

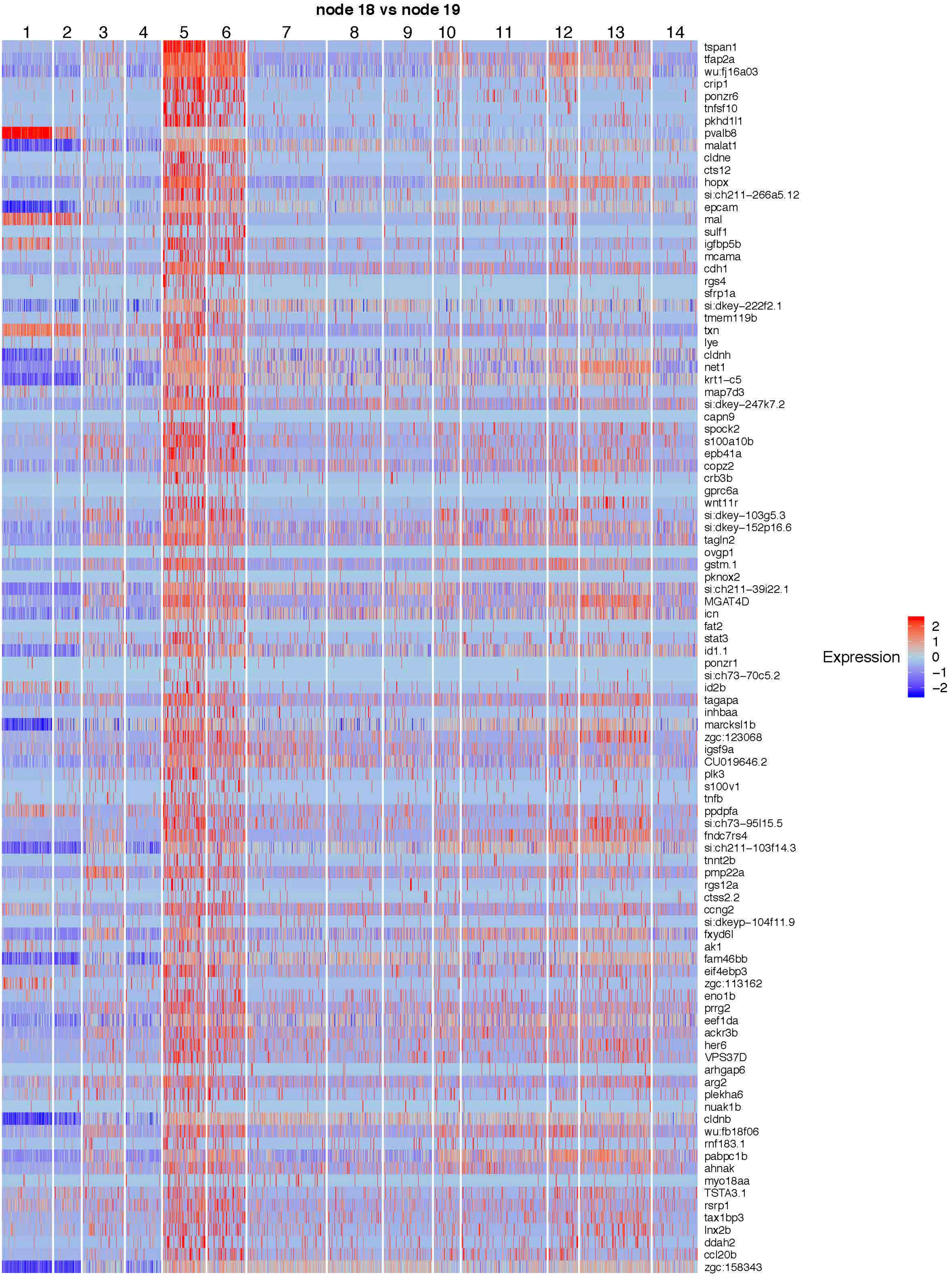
node 18 vs node 19
1
2
3
4
5
6
7
8
9
10
11
12
13
14
tspan1
tfap2a
wu:fj16a03
crip1
ponzr6
tnfsf10
pkhd1l1
pvalb8
malat1
cldne
cts12
hopx
si:ch211-266a5.12
epcam
mal
sulf1
igfbp5b
mcama
cdh1
rgs4
sfrp1a
si:dkey-222f2.1
tmem119b
txn
lye
cldnh
net1
krt1-c5
map7d3
si:dkey-247k7.2
capn9
spock2
s100a10b
epb41a
copz2
crb3b
gprc6a
wnt11r
si:dkey-103g5.3
si:dkey-152p16.6
tagln2
ovgp1
gstm.1
pknox2
si:ch211-39i22.1
MGAT4D
icn
fat2
stat3
id1.1
ponzr1
si:ch73-70c5.2
id2b
tagapa
inhbaa
marcksl1b
zgc:123068
igsf9a
CU019646.2
plk3
s100v1
tnfb
ppdpfa
si:ch73-95l15.5
fndc7rs4
si:ch211-103f14.3
tnnt2b
pmp22a
rgs12a
ctss2.2
ccng2
si:dkeyp-104f11.9
fxyd6l
ak1
fam46bb
eif4ebp3
zgc:113162
eno1b
prrg2
eef1da
ackr3b
her6
VPS37D
arhgap6
arg2
plekha6
nuak1b
cldnb
wu:fb18f06
rnf183.1
pabpc1b
ahnak
myo18aa
TSTA3.1
rsrp1
tax1bp3
lnx2b
ddah2
ccl20b
zgc:158343
Expression
2
1
0
−1
−2

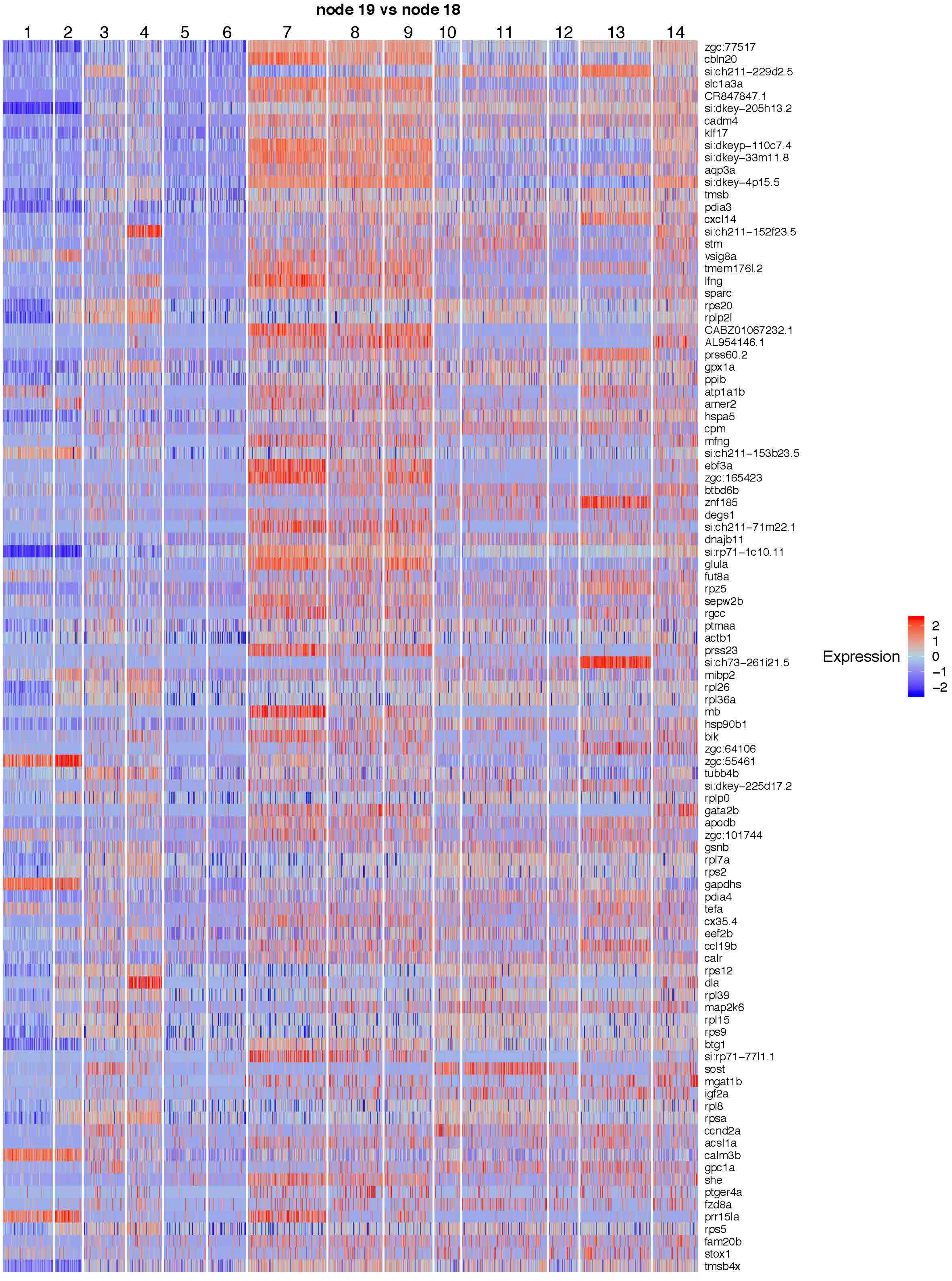
node 19 vs node 18
1
2
3
4
5
6
7
8
9
10
11
12
13
14
zgc:77517
cbln20
si:ch211-229d2.5
slc1a3a
CR847847.1
si:dkey-205h13.2
cadm4
klf17
si:dkeyp-110c7.4
si:dkey-33m11.8
aqp3a
si:dkey-4p15.5
tmsb
pdia3
cxcl14
si:ch211-152f23.5
stm
vsig8a
tmem176l.2
lfng
sparc
rps20
rplp2l
CABZ01067232.1
AL954146.1
prss60.2
gpx1a
ppib
atp1a1b
amer2
hspa5
cpm
mfng
si:ch211-153b23.5
ebf3a
zgc:165423
btbd6b
znf185
degs1
si:ch211-71m22.1
dnajb11
si:rp71-1c10.11
glula
fut8a
rpz5
sepw2b
rgcc
ptmaa
actb1
prss23
si:ch73-261i21.5
mibp2
rpl26
rpl36a
mb
hsp90b1
bik
zgc:64106
zgc:55461
tubb4b
si:dkey-225d17.2
rplp0
gata2b
apodb
zgc:101744
gsnb
rpl7a
rps2
gapdhs
pdia4
tefa
cx35.4
eef2b
ccl19b
calr
rps12
dla
rpl39
map2k6
rpl15
rps9
btg1
si:rp71-77l1.1
sost
mgat1b
igf2a
rpl8
rpsa
ccnd2a
acsl1a
calm3b
gpc1a
she
ptger4a
fzd8a
prr15la
rps5
fam20b
stox1
tmsb4x
Expression
2
1
0
−1
−2

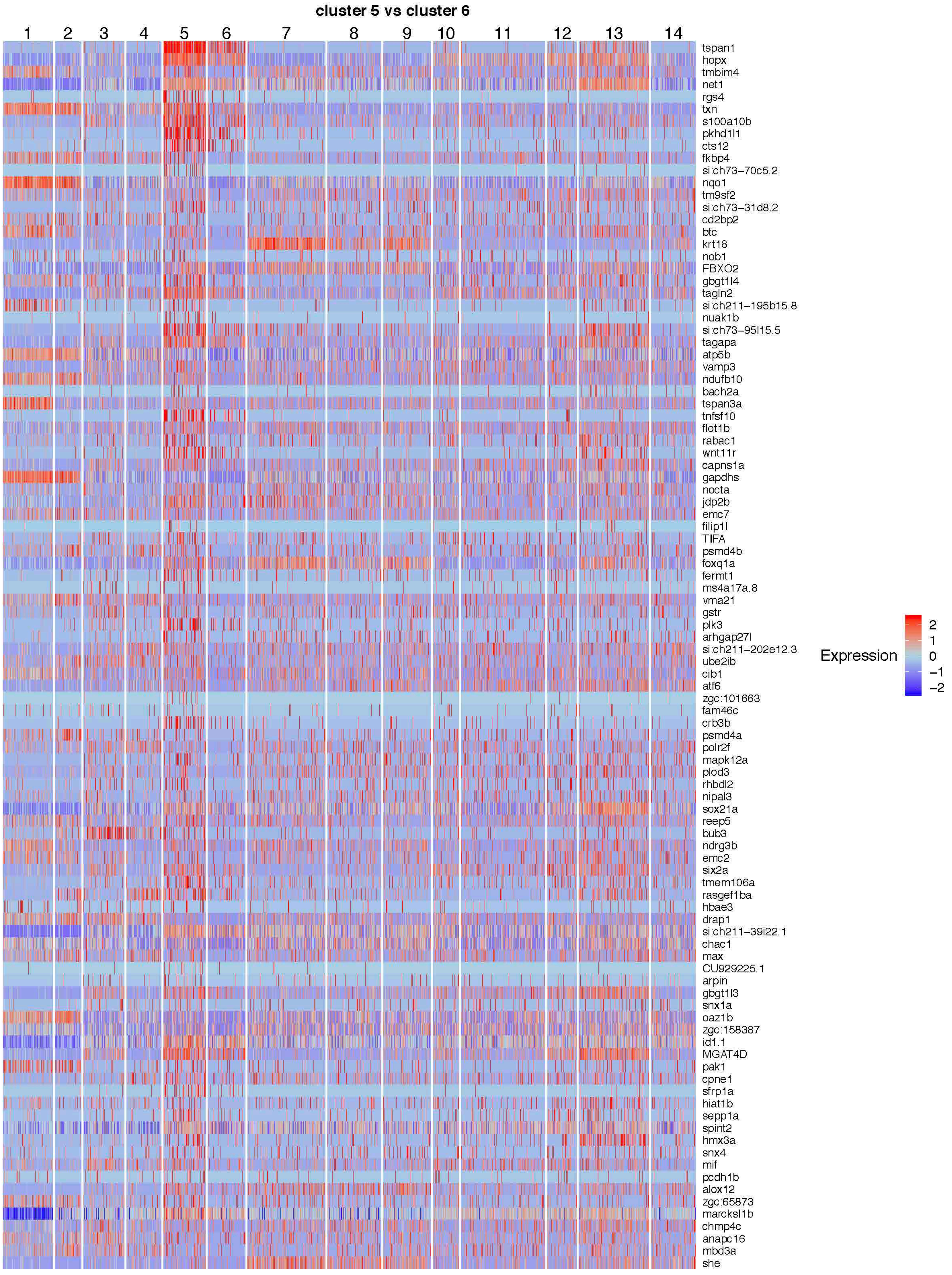
cluster 5 vs cluster 6
1
2
3
4
5
6
7
8
9
10
11
12
13
14
tspan1
hopx
tmbim4
net1
rgs4
txn
s100a10b
pkhd1l1
cts12
fkbp4
si:ch73-70c5.2
nqo1
tm9sf2
si:ch73-31d8.2
cd2bp2
btc
krt18
nob1
FBXO2
gbgt1l4
tagln2
si:ch211-195b15.8
nuak1b
si:ch73-95l15.5
tagapa
atp5b
vamp3
ndufb10
bach2a
tspan3a
tnfsf10
flot1b
rabac1
wnt11r
capns1a
gapdhs
nocta
jdp2b
emc7
filip1l
TIFA
psmd4b
foxq1a
fermt1
ms4a17a.8
vma21
gstr
plk3
arhgap27l
si:ch211-202e12.3
ube2ib
cib1
atf6
zgc:101663
fam46c
crb3b
psmd4a
polr2f
mapk12a
plod3
rhbdl2
nipal3
sox21a
reep5
bub3
ndrg3b
emc2
six2a
tmem106a
rasgef1ba
hbae3
drap1
si:ch211-39i22.1
chac1
max
CU929225.1
arpin
gbgt1l3
snx1a
oaz1b
zgc:158387
id1.1
MGAT4D
pak1
cpne1
sfrp1a
hiat1b
sepp1a
spint2
hmx3a
snx4
mif
pcdh1b
alox12
zgc:65873
marcksl1b
chmp4c
anapc16
mbd3a
she
Expression
2
1
0
−1
−2

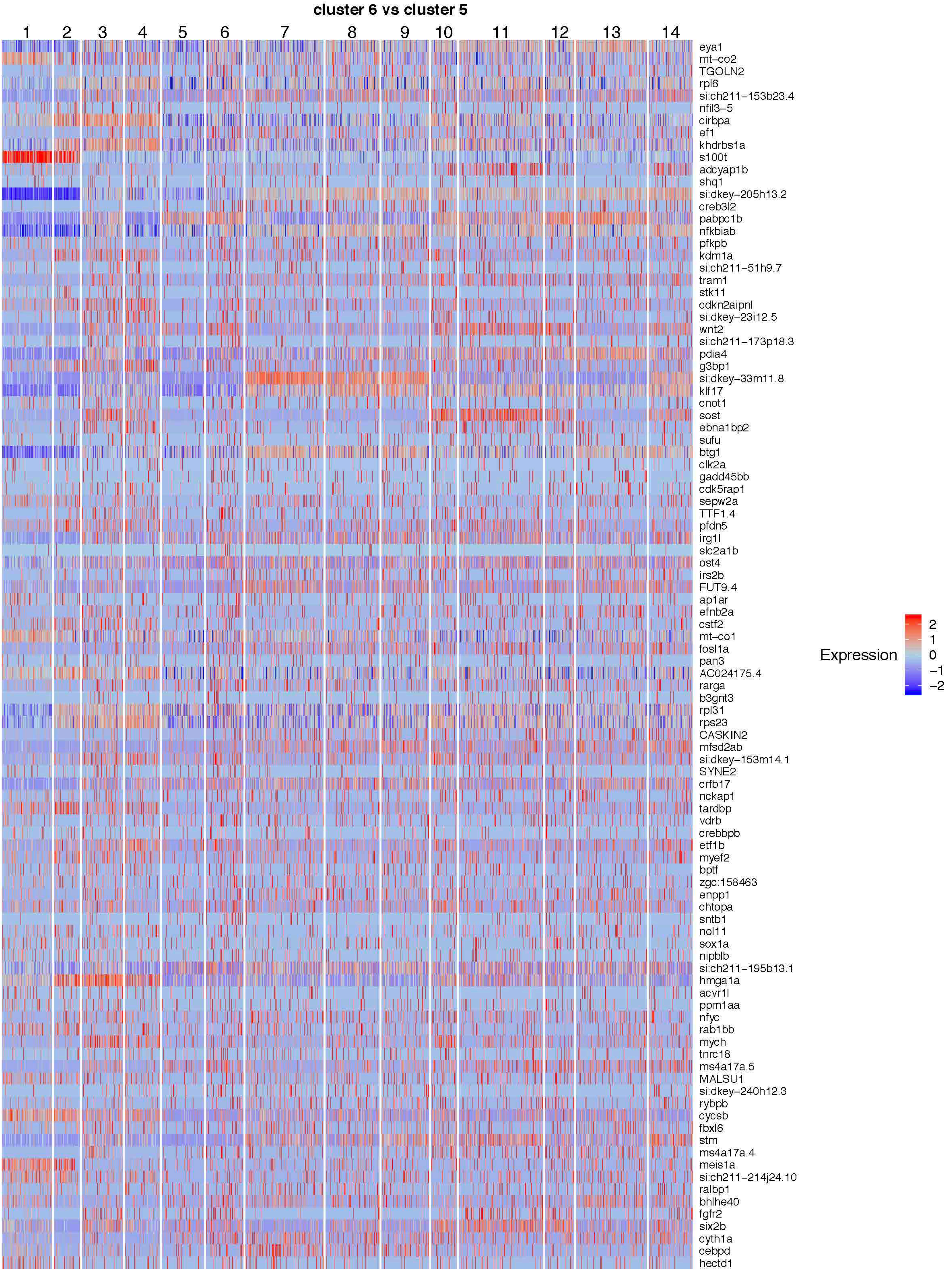
cluster 6 vs cluster 5
1
2
3
4
5
6
7
8
9
10
11
12
13
14
eya1
mt−co2
TGOLN2
rpl6
si:ch211−153b23.4
nfil3−5
cirbpa
ef1
khdrbs1a
s100t
adcyap1b
shq1
si:dkey−205h13.2
creb3l2
pabpc1b
nfkbiab
pfkpb
kdm1a
si:ch211−51h9.7
tram1
stk11
cdkn2aipnl
si:dkey−23i12.5
wnt2
si:ch211−173p18.3
pdia4
g3bp1
si:dkey−33m11.8
klf17
cnot1
sost
ebna1bp2
sufu
btg1
clk2a
gadd45bb
cdk5rap1
sepw2a
TTF1.4
pfdn5
irg1l
slc2a1b
ost4
irs2b
FUT9.4
ap1ar
efnb2a
cstf2
mt−co1
fosl1a
pan3
AC024175.4
rarga
b3gnt3
rpl31
rps23
CASKIN2
mfsd2ab
si:dkey−153m14.1
SYNE2
crfb17
nckap1
tardbp
vdrb
crebbpb
etf1b
myef2
bptf
zgc:158463
enpp1
chtopa
sntb1
nol11
sox1a
nipblb
si:ch211−195b13.1
hmga1a
acvr1l
ppm1aa
nfyc
rab1bb
mych
tnrc18
ms4a17a.5
MALSU1
si:dkey−240h12.3
rybpb
cycsb
fbxl6
stm
ms4a17a.4
meis1a
si:ch211−214j24.10
ralbp1
bhlhe40
fgfr2
six2b
cyth1a
cebpd
hectd1
Expression
2
1
0
−1
−2

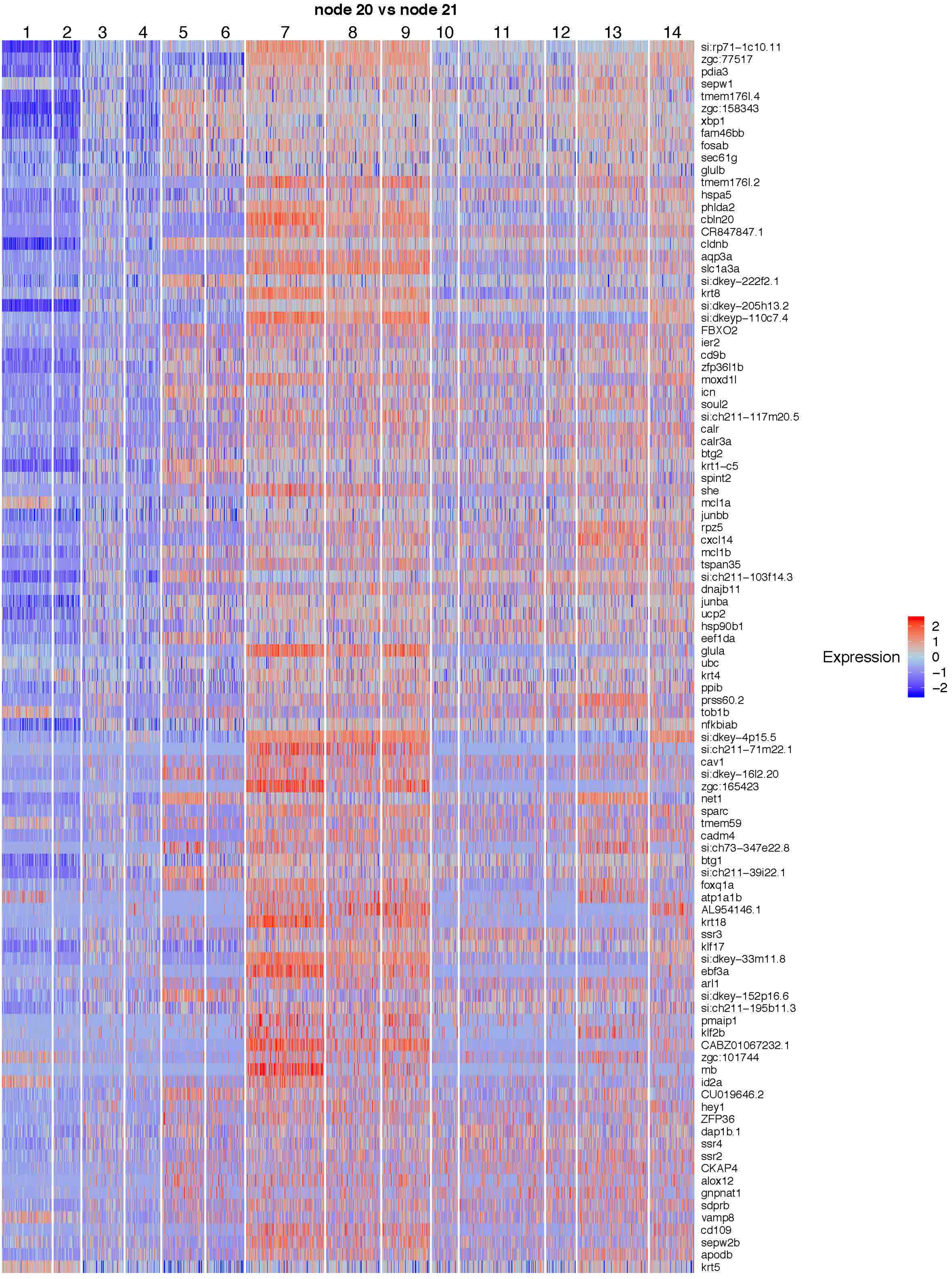
node 20 vs node 21
1
2
3
4
5
6
7
8
9
10
11
12
13
14
si:rp71−1c10.11
zgc:77517
pdia3
sepw1
tmem176l.4
zgc:158343
xbp1
fam46bb
fosab
sec61g
glulb
tmem176l.2
hspa5
phlda2
cbln20
CR847847.1
cldnb
aqp3a
slc1a3a
si:dkey−222f2.1
krt8
si:dkey−205h13.2
si:dkeyp−110c7.4
FBXO2
ier2
cd9b
zfp36l1b
moxd1l
icn
soul2
si:ch211−117m20.5
calr
calr3a
btg2
krt1−c5
spint2
she
mcl1a
junbb
rpz5
cxcl14
mcl1b
tspan35
si:ch211−103f14.3
dnajb11
junba
ucp2
hsp90b1
eef1da
glula
ubc
krt4
ppib
prss60.2
tob1b
nfkbiab
si:dkey−4p15.5
si:ch211−71m22.1
cav1
si:dkey−16l2.20
zgc:165423
net1
sparc
tmem59
cadm4
si:ch73−347e22.8
btg1
si:ch211−39i22.1
foxq1a
atp1a1b
AL954146.1
krt18
ssr3
klf17
si:dkey−33m11.8
ebf3a
arl1
si:dkey−152p16.6
si:ch211−195b11.3
pmaip1
klf2b
CABZ01067232.1
zgc:101744
mb
id2a
CU019646.2
hey1
ZFP36
dap1b.1
ssr4
ssr2
CKAP4
alox12
gnpnat1
sdprb
vamp8
cd109
sepw2b
apodb
krt5
Expression
2
1
0
−1
−2

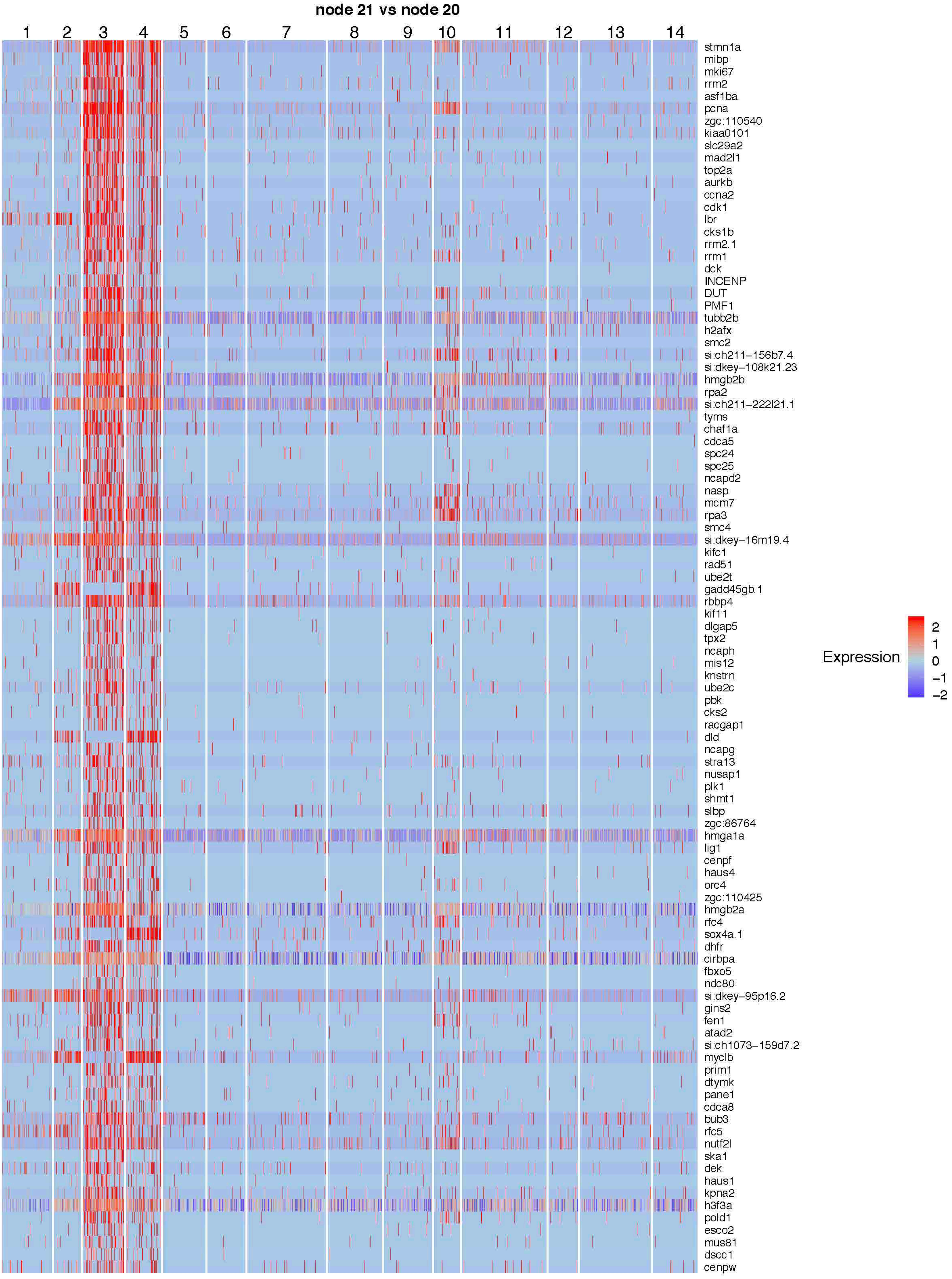
node 21 vs node 20
1
2
3
4
5
6
7
8
9
10
11
12
13
14
stmn1a
mibp
mki67
rrm2
asf1ba
pcna
zgc:110540
kiaa0101
slc29a2
mad2l1
top2a
aurkb
ccna2
cdk1
lbr
cks1b
rrm2.1
rrm1
dck
INCENP
DUT
PMF1
tubb2b
h2afx
smc2
si:ch211−156b7.4
si:dkey−108k21.23
hmgb2b
rpa2
si:ch211−222l21.1
tyms
chaf1a
cdca5
spc24
spc25
ncapd2
nasp
mcm7
rpa3
smc4
si:dkey−16m19.4
kifc1
rad51
ube2t
gadd45gb.1
rbbp4
kif11
dlgap5
tpx2
ncaph
mis12
knstrn
ube2c
pbk
cks2
racgap1
dld
ncapg
stra13
nusap1
plk1
shmt1
slbp
zgc:86764
hmga1a
lig1
cenpf
haus4
orc4
zgc:110425
hmgb2a
rfc4
sox4a.1
dhfr
cirbpa
fbxo5
ndc80
si:dkey−95p16.2
gins2
fen1
atad2
si:ch1073−159d7.2
myclb
prim1
dtymk
pane1
cdca8
bub3
rfc5
nutf2l
ska1
dek
haus1
kpna2
h3f3a
pold1
esco2
mus81
dscc1
cenpw
Expression
2
1
0
−1
−2

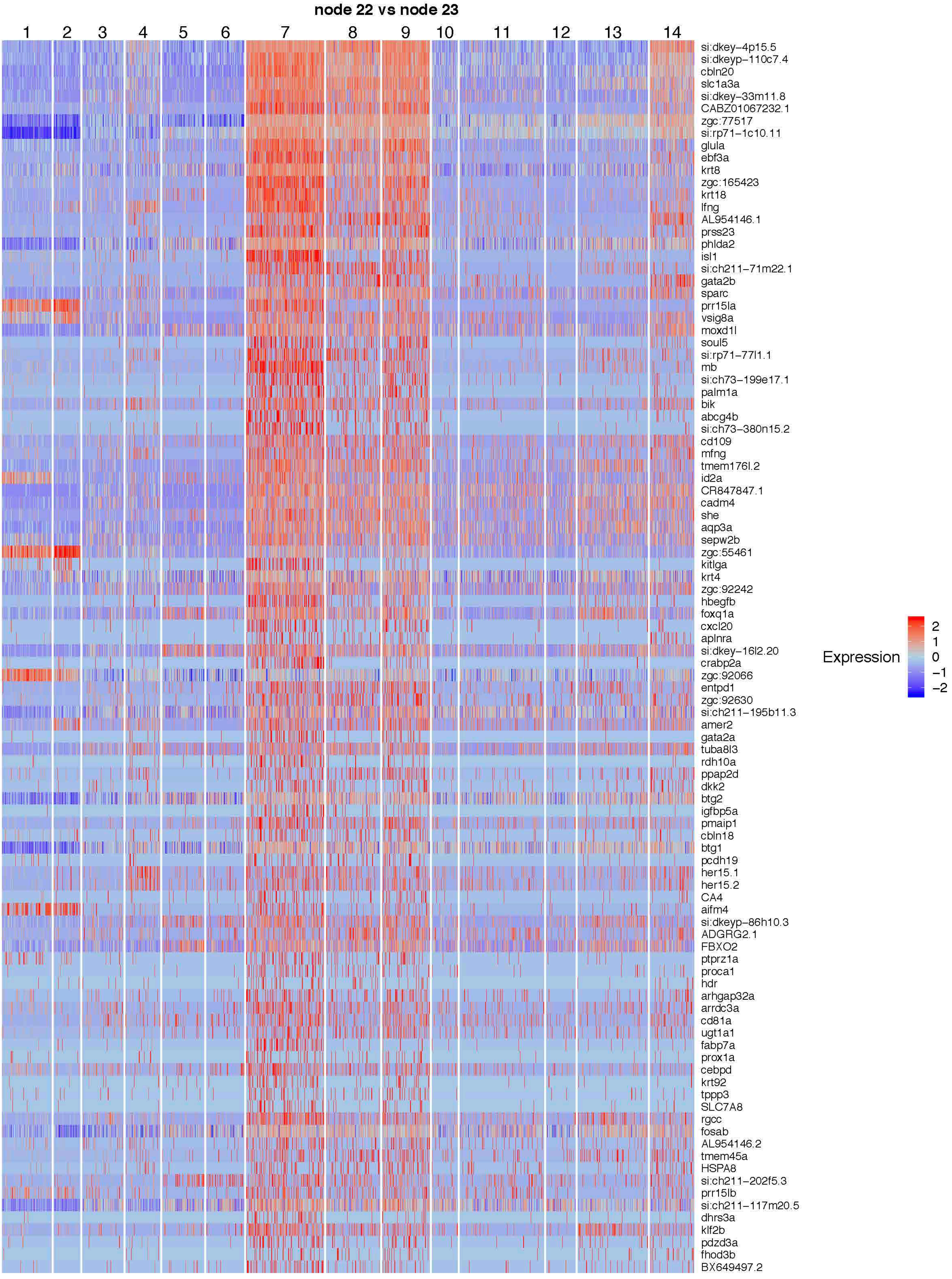
node 22 vs node 23
1
2
3
4
5
6
7
8
9
10
11
12
13
14
si:dkey−4p15.5
si:dkeyp−110c7.4
cbln20
slc1a3a
si:dkey−33m11.8
CABZ01067232.1
zgc:77517
si:rp71−1c10.11
glula
ebf3a
krt8
zgc:165423
krt18
lfng
AL954146.1
prss23
phlda2
isl1
si:ch211−71m22.1
gata2b
sparc
prr15la
vsig8a
moxd1l
soul5
si:rp71−77l1.1
mb
si:ch73−199e17.1
palm1a
bik
abcg4b
si:ch73−380n15.2
cd109
mfng
tmem176l.2
id2a
CR847847.1
cadm4
she
aqp3a
sepw2b
zgc:55461
kitlga
krt4
zgc:92242
hbegfb
foxq1a
cxcl20
aplnra
si:dkey−16l2.20
crabp2a
zgc:92066
entpd1
zgc:92630
si:ch211−195b11.3
amer2
gata2a
tuba8l3
rdh10a
ppap2d
dkk2
btg2
igfbp5a
pmaip1
cbln18
btg1
pcdh19
her15.1
her15.2
CA4
aifm4
si:dkeyp−86h10.3
ADGRG2.1
FBXO2
ptprz1a
proca1
hdr
arhgap32a
arrdc3a
cd81a
ugt1a1
fabp7a
prox1a
cebpd
krt92
tppp3
SLC7A8
rgcc
fosab
AL954146.2
tmem45a
HSPA8
si:ch211−202f5.3
prr15lb
si:ch211−117m20.5
dhrs3a
klf2b
pdzd3a
fhod3b
BX649497.2
Expression
2
1
0
−1
−2

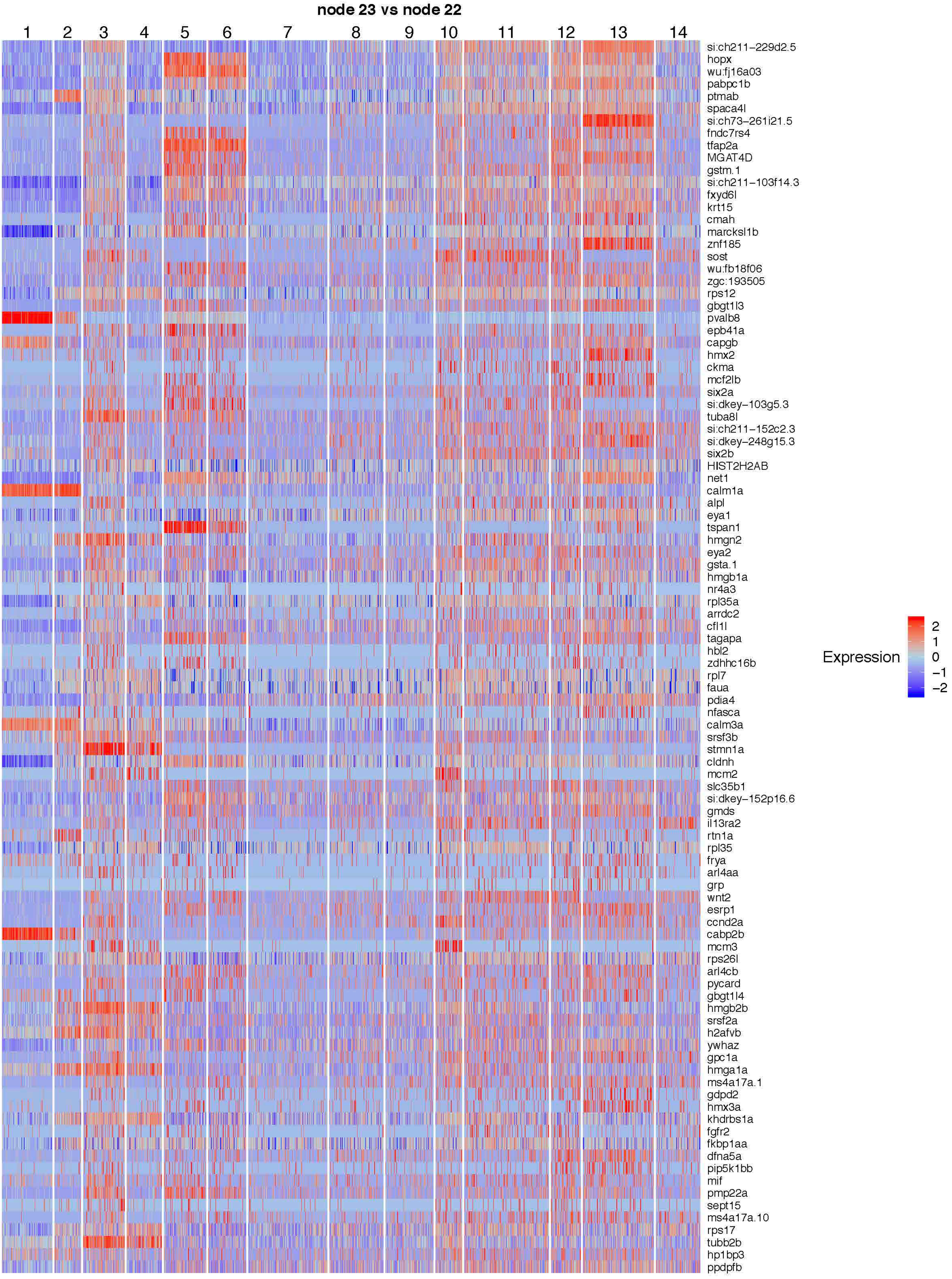

node 23 vs node 22
1
2
3
4
5
6
7
8
9
10
11
12
13
14
si:ch211−229d2.5
hopx
wu:fj16a03
pabpc1b
ptmab
spaca4l
si:ch73−261i21.5
fndc7rs4
tfap2a
MGAT4D
gstm.1
si:ch211−103f14.3
fxyd6l
krt15
cmah
marcksl1b
znf185
sost
wu:fb18f06
zgc:193505
rps12
gbgt1l3
pvalb8
epb41a
capgb
hmx2
ckma
mcf2lb
six2a
si:dkey−103g5.3
tuba8l
si:ch211−152c2.3
si:dkey−248g15.3
six2b
HIST2H2AB
net1
calm1a
alpl
eya1
tspan1
hmgn2
eya2
gsta.1
hmgb1a
nr4a3
rpl35a
arrdc2
cfl1l
tagapa
hbl2
zdhhc16b
rpl7
faua
pdia4
nfasca
calm3a
srsf3b
stmn1a
cldnh
mcm2
slc35b1
si:dkey−152p16.6
gmds
il13ra2
rtn1a
rpl35
frya
arl4aa
grp
wnt2
esrp1
ccnd2a
cabp2b
mcm3
rps26l
arl4cb
pycard
gbgt1l4
hmgb2b
srsf2a
h2afvb
ywhaz
gpc1a
hmga1a
ms4a17a.1
gdpd2
hmx3a
khdrbs1a
fgfr2
fkbp1aa
dfna5a
pip5k1bb
mif
pmp22a
sept15
ms4a17a.10
rps17
tubb2b
hp1bp3
ppdpfb
Expression
2
1
0
−1
−2

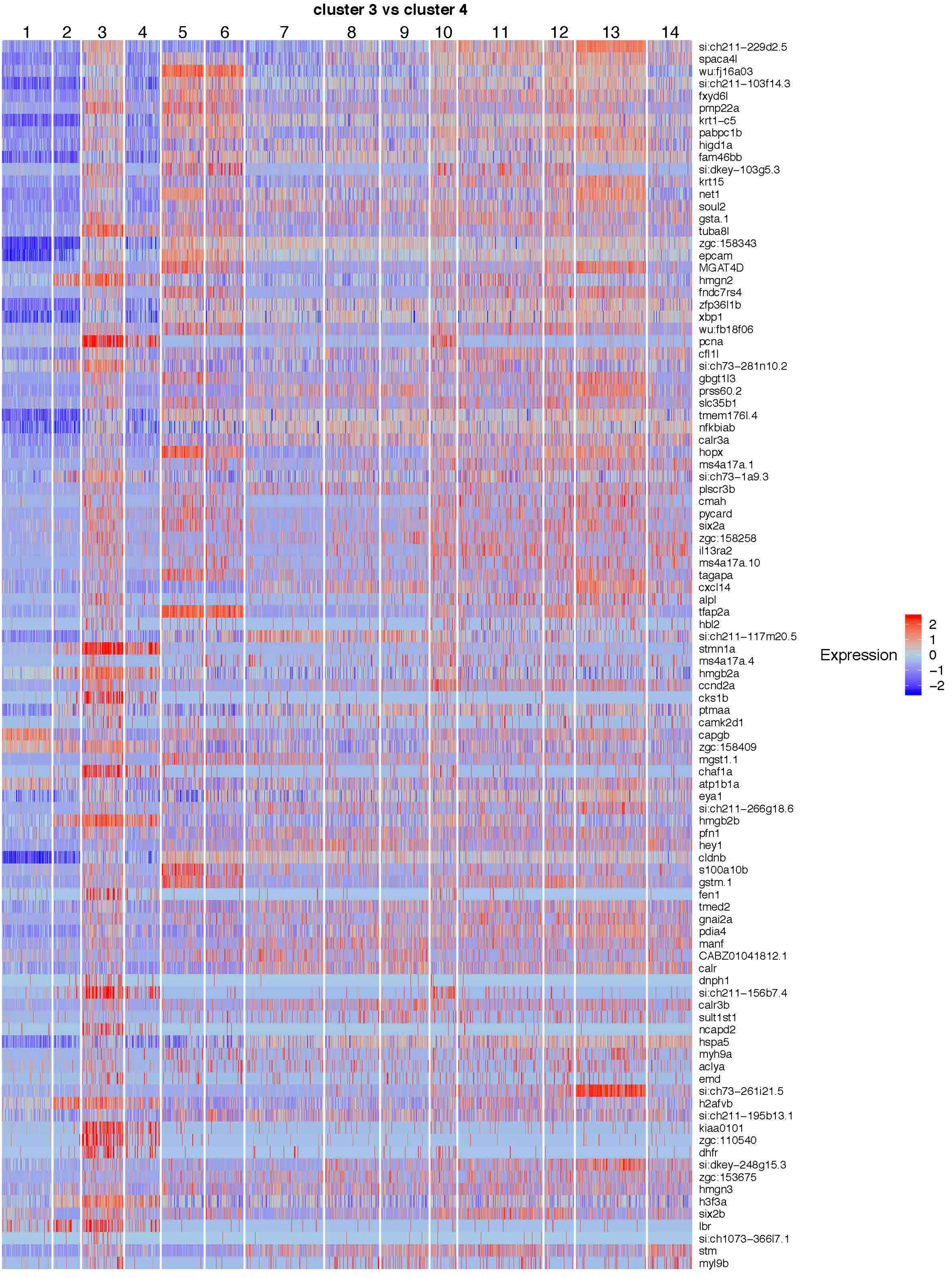
cluster 3 vs cluster 4
1
2
3
4
5
6
7
8
9
10
11
12
13
14
si:ch211-229d2.5
spaca4l
wu:fj16a03
si:ch211-103f14.3
fxyd6l
pmp22a
krt1-c5
pabpc1b
higd1a
fam46bb
si:dkey-103g5.3
krt15
net1
soul2
gsta.1
tuba8l
zgc:158343
epcam
MGAT4D
hmgn2
fndc7rs4
zfp36l1b
xbp1
wu:fb18f06
pcna
cfl1l
si:ch73-281n10.2
gbgt1l3
prss60.2
slc35b1
tmem176l.4
nfkbiab
calr3a
hopx
ms4a17a.1
si:ch73-1a9.3
plscr3b
cmah
pycard
six2a
zgc:158258
il13ra2
ms4a17a.10
tagapa
cxcl14
alpl
tfap2a
hbl2
si:ch211-117m20.5
stmn1a
ms4a17a.4
hmgb2a
ccnd2a
cks1b
ptmaa
camk2d1
capgb
zgc:158409
mgst1.1
chaf1a
atp1b1a
eya1
si:ch211-266g18.6
hmgb2b
pfn1
hey1
cldnb
s100a10b
gstm.1
fen1
tmed2
gnai2a
pdia4
manf
CABZ01041812.1
calr
dnph1
si:ch211-156b7.4
calr3b
sult1st1
ncapd2
hspa5
myh9a
aclya
emd
si:ch73-261i21.5
h2afvb
si:ch211-195b13.1
kiaa0101
zgc:110540
dhfr
si:dkey-248g15.3
zgc:153675
hmgn3
h3f3a
six2b
lbr
si:ch1073-366l7.1
stm
myl9b
Expression
2
1
0
−1
−2

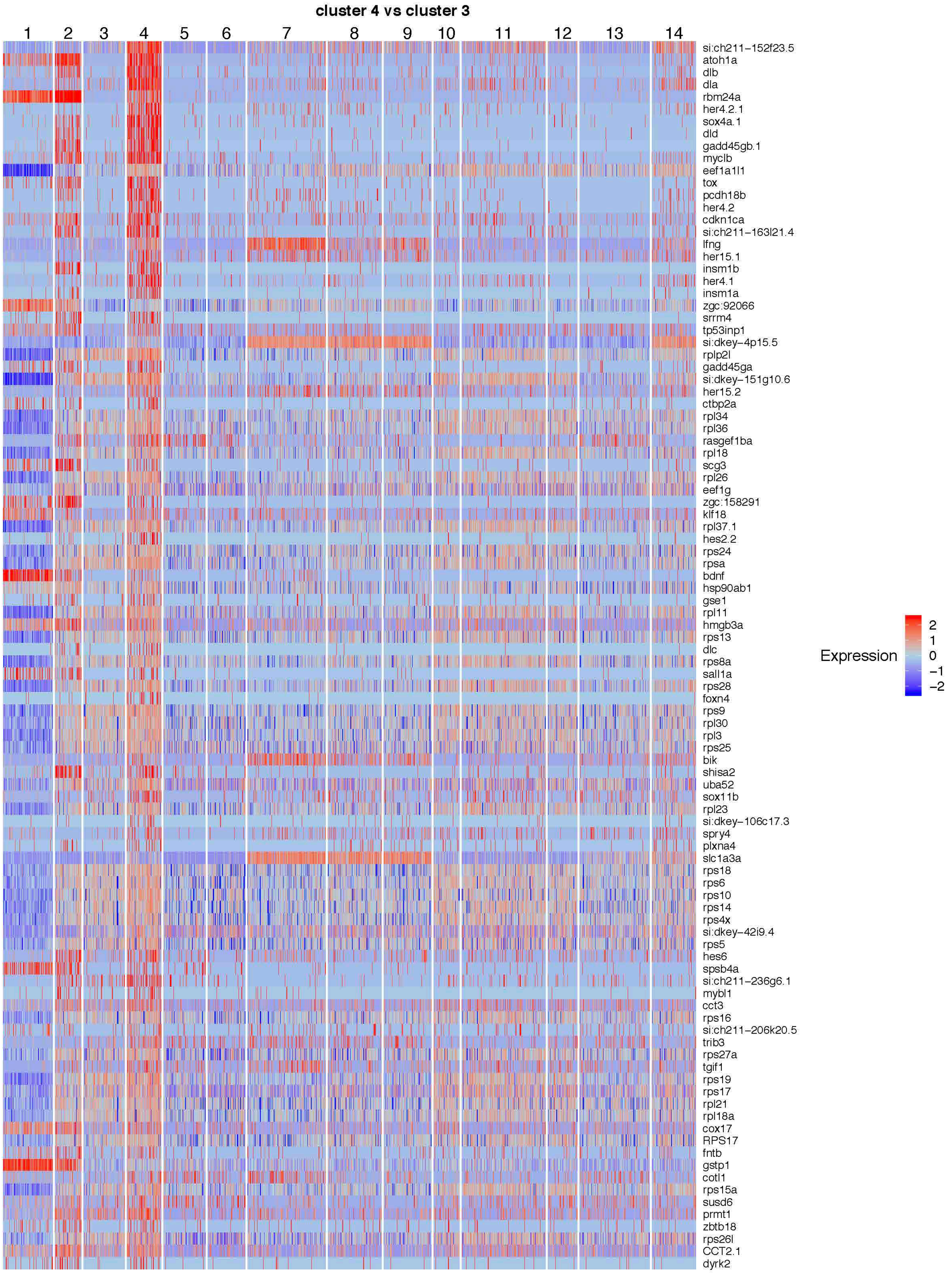
cluster 4 vs cluster 3
1
2
3
4
5
6
7
8
9
10
11
12
13
14
si:ch211−152f23.5
atoh1a
dlb
dla
rbm24a
her4.2.1
sox4a.1
dld
gadd45gb.1
myclb
eef1a1l1
tox
pcdh18b
her4.2
cdkn1ca
si:ch211−163l21.4
lfng
her15.1
insm1b
her4.1
insm1a
zgc:92066
srrm4
tp53inp1
si:dkey−4p15.5
rplp2l
gadd45ga
si:dkey−151g10.6
her15.2
ctbp2a
rpl34
rpl36
rasgef1ba
rpl18
scg3
rpl26
eef1g
zgc:158291
klf18
rpl37.1
hes2.2
rps24
rpsa
bdnf
hsp90ab1
gse1
rpl11
hmgb3a
rps13
dlc
rps8a
sall1a
rps28
foxn4
rps9
rpl30
rpl3
rps25
bik
shisa2
uba52
sox11b
rpl23
si:dkey−106c17.3
spry4
plxna4
slc1a3a
rps18
rps6
rps10
rps14
rps4x
si:dkey−42i9.4
rps5
hes6
spsb4a
si:ch211−236g6.1
mybl1
cct3
rps16
si:ch211−206k20.5
trib3
rps27a
tgif1
rps19
rps17
rpl21
rpl18a
cox17
RPS17
fntb
gstp1
cotl1
rps15a
susd6
prmt1
zbtb18
rps26l
CCT2.1
dyrk2
Expression
2
1
0
−1
−2

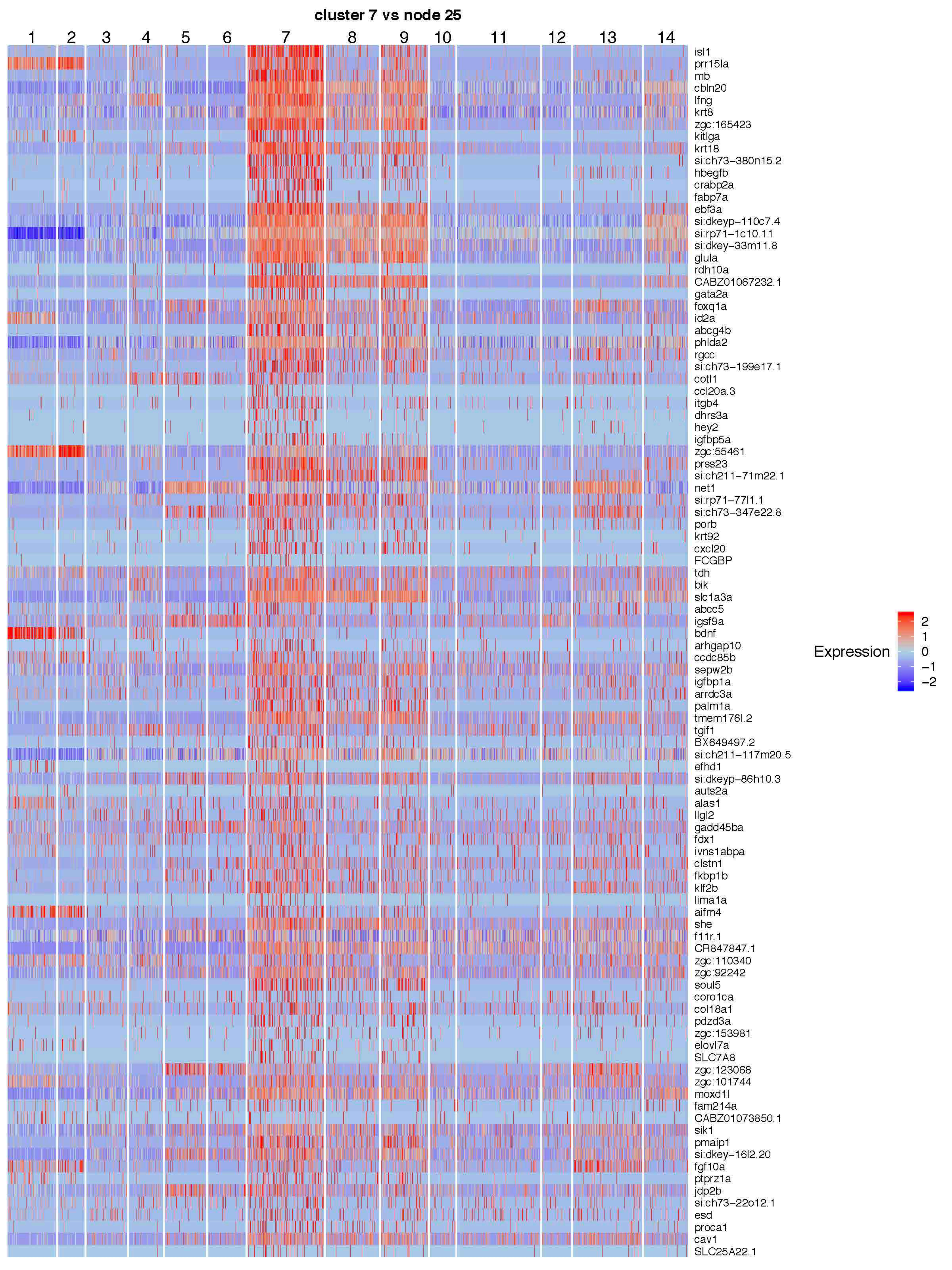
cluster 7 vs node 25
1
2
3
4
5
6
7
8
9
10
11
12
13
14
isl1
prr15la
mb
cbln20
lfng
krt8
zgc:165423
kitlga
krt18
si:ch73−380n15.2
hbegfb
crabp2a
fabp7a
ebf3a
si:dkeyp−110c7.4
si:rp71−1c10.11
si:dkey−33m11.8
glula
rdh10a
CABZ01067232.1
gata2a
foxq1a
id2a
abcg4b
phlda2
rgcc
si:ch73−199e17.1
cotl1
ccl20a.3
itgb4
dhrs3a
hey2
igfbp5a
zgc:55461
prss23
si:ch211−71m22.1
net1
si:rp71−77l1.1
si:ch73−347e22.8
porb
krt92
cxcl20
FCGBP
tdh
bik
slc1a3a
abcc5
igsf9a
bdnf
arhgap10
ccdc85b
sepw2b
igfbp1a
arrdc3a
palm1a
tmem176l.2
tgif1
BX649497.2
si:ch211−117m20.5
efhd1
si:dkeyp−86h10.3
auts2a
alas1
llgl2
gadd45ba
fdx1
ivns1abpa
clstn1
fkbp1b
klf2b
lima1a
aifm4
she
f11r.1
CR847847.1
zgc:110340
zgc:92242
soul5
coro1ca
col18a1
pdzd3a
zgc:153981
elovl7a
SLC7A8
zgc:123068
zgc:101744
moxd1l
fam214a
CABZ01073850.1
sik1
pmaip1
si:dkey−16l2.20
fgf10a
ptprz1a
jdp2b
si:ch73−22o12.1
esd
proca1
cav1
SLC25A22.1
Expression
2
1
0
−1
−2

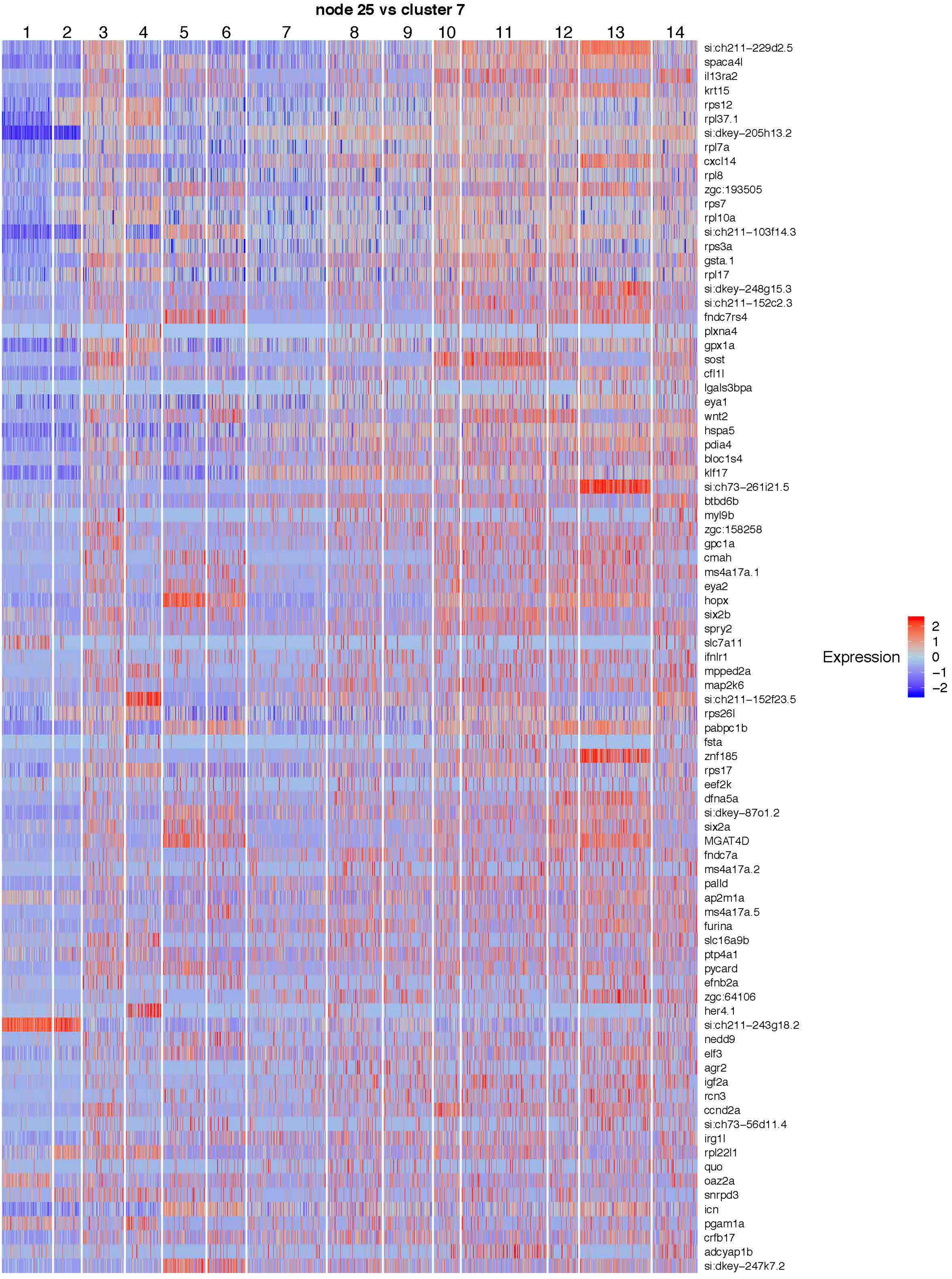
node 25 vs cluster 7
1
2
3
4
5
6
7
8
9
10
11
12
13
14
si:ch211-229d2.5
spaca4l
il13ra2
krt15
rps12
rpl37.1
si:dkey-205h13.2
rpl7a
cxcl14
rpl8
zgc:193505
rps7
rpl10a
si:ch211-103f14.3
rps3a
gsta.1
rpl17
si:dkey-248g15.3
si:ch211-152c2.3
fndc7rs4
plxna4
gpx1a
sost
cfl1l
lgals3bpa
eya1
wnt2
hspa5
pdia4
bloc1s4
klf17
si:ch73-261i21.5
btbd6b
myl9b
zgc:158258
gpc1a
cmah
ms4a17a.1
eya2
hopx
six2b
spry2
slc7a11
ifnlr1
mpped2a
map2k6
si:ch211-152f23.5
rps26l
pabpc1b
fsta
znf185
rps17
eef2k
dfna5a
si:dkey-87o1.2
six2a
MGAT4D
fndc7a
ms4a17a.2
palld
ap2m1a
ms4a17a.5
furina
slc16a9b
ptp4a1
pycard
efnb2a
zgc:64106
her4.1
si:ch211-243g18.2
nedd9
elf3
agr2
igf2a
rcn3
ccnd2a
si:ch73-56d11.4
irg1l
rpl22l1
quo
oaz2a
snrpd3
icn
pgam1a
crfb17
adcyap1b
si:dkey-247k7.2
Expression
2
1
0
−1
−2

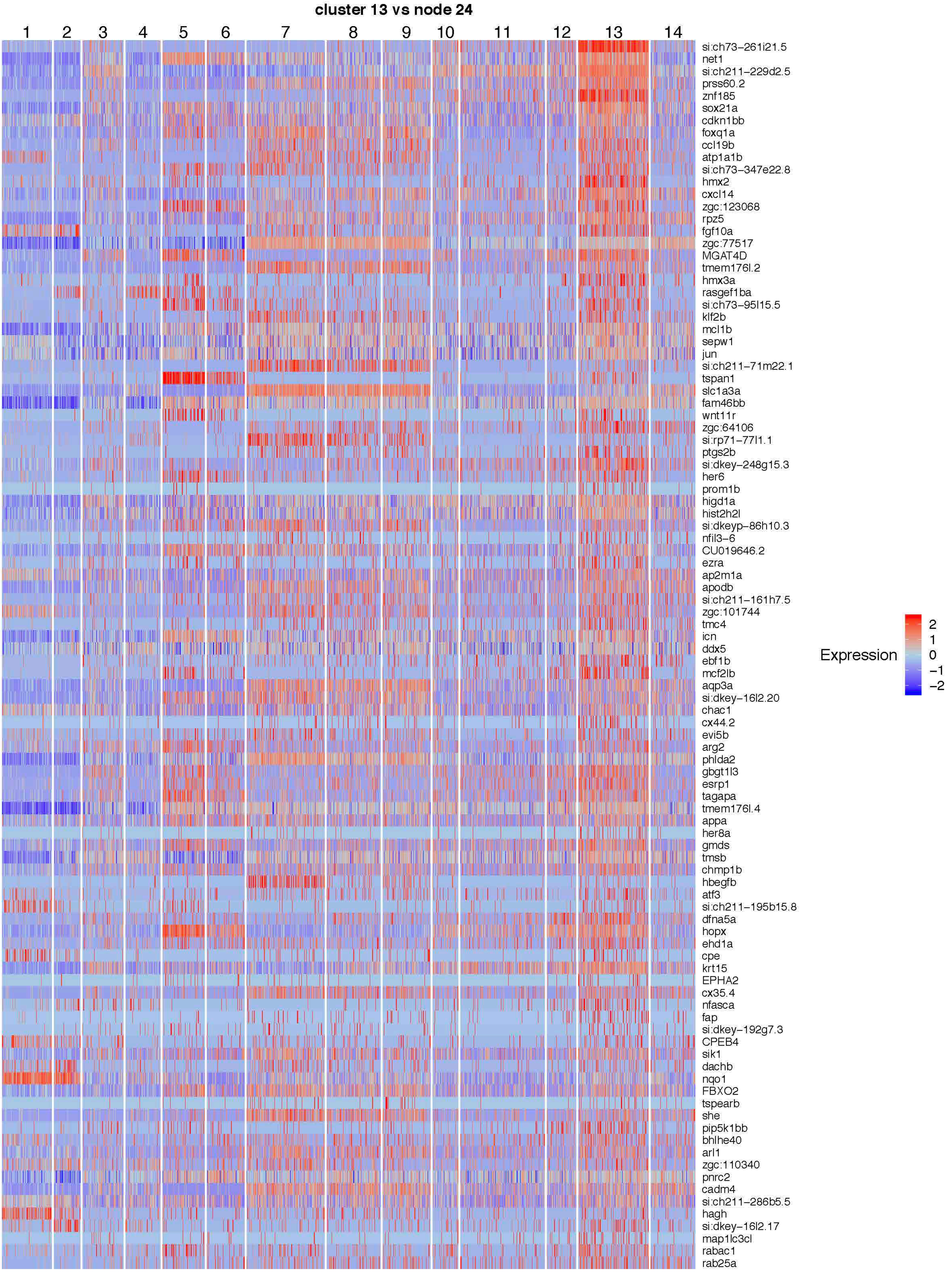
cluster 13 vs node 24
1
2
3
4
5
6
7
8
9
10
11
12
13
14
si:ch73-261i21.5
net1
si:ch211-229d2.5
prss60.2
znf185
sox21a
cdkn1bb
foxq1a
ccl19b
atp1a1b
si:ch73-347e22.8
hmx2
cxcl14
zgc:123068
rpz5
fgf10a
zgc:77517
MGAT4D
tmem176l.2
hmx3a
rasgef1ba
si:ch73-95l15.5
klf2b
mcl1b
sepw1
jun
si:ch211-71m22.1
tspan1
slc1a3a
fam46bb
wnt11r
zgc:64106
si:rp71-77l1.1
ptgs2b
si:dkey-248g15.3
her6
prom1b
higd1a
hist2h2l
si:dkeyp-86h10.3
nfil3-6
CU019646.2
ezra
ap2m1a
apodb
si:ch211-161h7.5
zgc:101744
tmc4
icn
ddx5
ebf1b
mcf2lb
aqp3a
si:dkey-16l2.20
chac1
cx44.2
evi5b
arg2
phlda2
gbgt1l3
esrp1
tagapa
tmem176l.4
appa
her8a
gmds
tmsb
chmp1b
hbegfb
atf3
si:ch211-195b15.8
dfna5a
hopx
ehd1a
cpe
krt15
EPHA2
cx35.4
nfasca
fap
si:dkey-192g7.3
CPEB4
sik1
dachb
nqo1
FBXO2
tspearb
she
pip5k1bb
bhlhe40
arl1
zgc:110340
pnrc2
cadm4
si:ch211-286b5.5
hagh
si:dkey-16l2.17
map1lc3cl
rabac1
rab25a
Expression
2
1
0
−1
−2

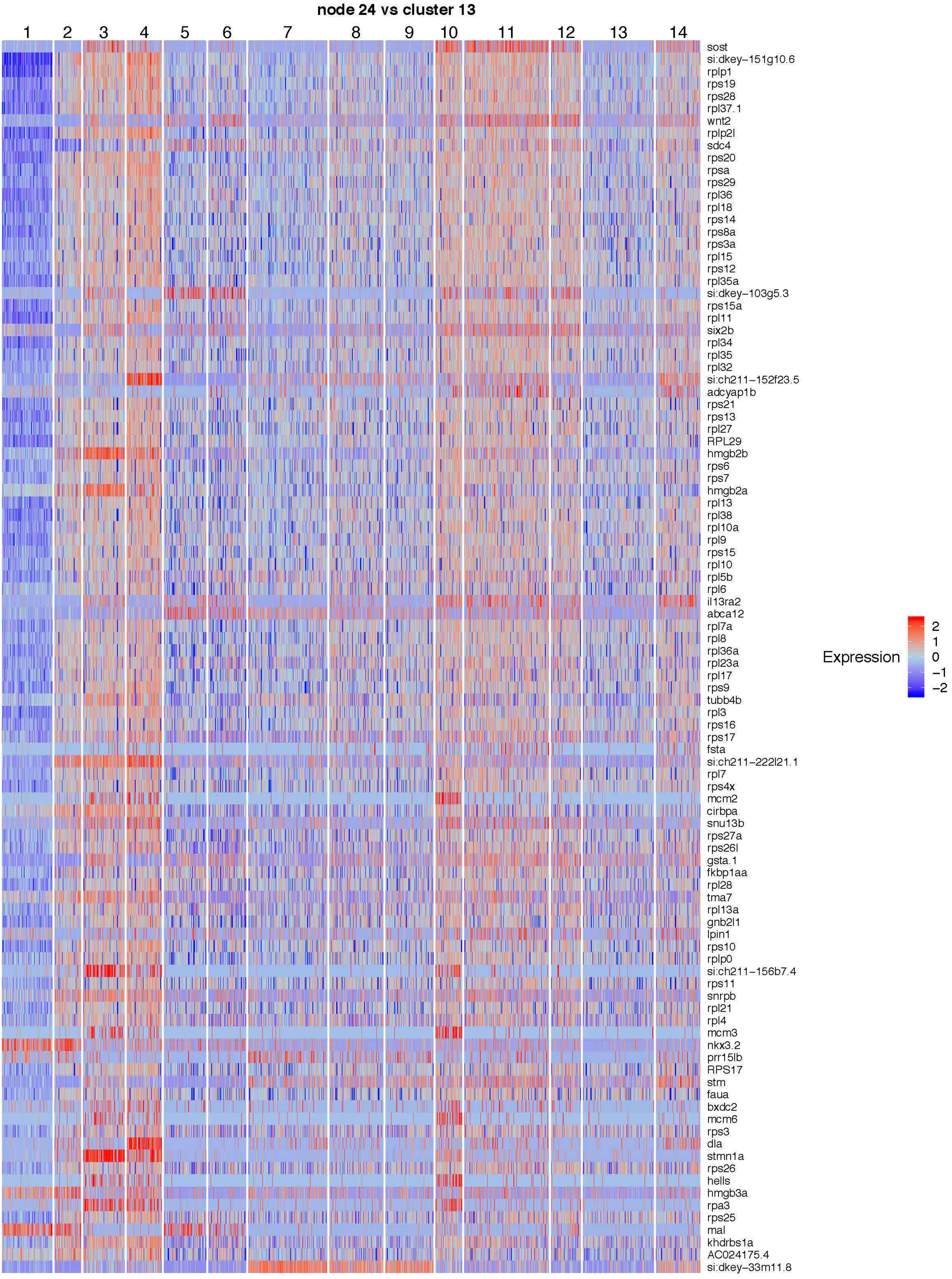

node 24 vs cluster 13
1
2
3
4
5
6
7
8
9
10
11
12
13
14
sost
si:dkey−151g10.6
rplp1
rps19
rps28
rpl37.1
wnt2
rplp2l
sdc4
rps20
rpsa
rps29
rpl36
rpl18
rps14
rps8a
rps3a
rpl15
rps12
rpl35a
si:dkey−103g5.3
rps15a
rpl11
six2b
rpl34
rpl35
rpl32
si:ch211−152f23.5
adcyap1b
rps21
rps13
rpl27
RPL29
hmgb2b
rps6
rps7
hmgb2a
rpl13
rpl38
rpl10a
rpl9
rps15
rpl10
rpl5b
rpl6
il13ra2
abca12
rpl7a
rpl8
rpl36a
rpl23a
rpl17
rps9
tubb4b
rpl3
rps16
rps17
fsta
si:ch211−222l21.1
rpl7
rps4x
mcm2
cirbpa
snu13b
rps27a
rps26l
gsta.1
fkbp1aa
rpl28
tma7
rpl13a
gnb2l1
lpin1
rps10
rplp0
si:ch211−156b7.4
rps11
snrpb
rpl21
rpl4
mcm3
nkx3.2
prr15lb
RPS17
stm
faua
bxdc2
mcm6
rps3
dla
stmn1a
rps26
hells
hmgb3a
rpa3
rps25
mal
khdrbs1a
AC024175.4
si:dkey−33m11.8
Expression
2
1
0
−1
−2

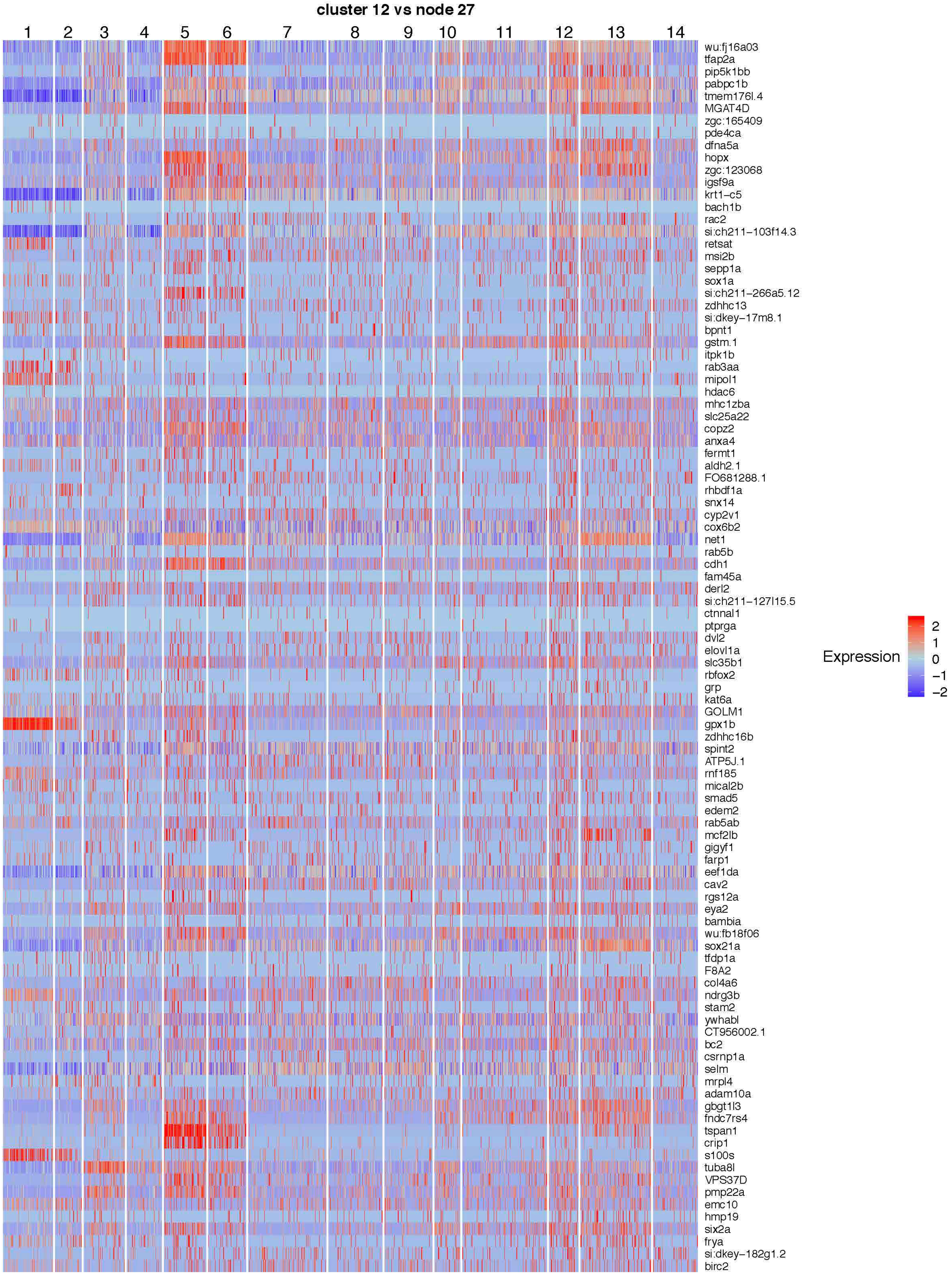
cluster 12 vs node 27
1
2
3
4
5
6
7
8
9
10
11
12
13
14
wu:fj16a03
tfap2a
pip5k1bb
pabpc1b
tmem176l.4
MGAT4D
zgc:165409
pde4ca
dfna5a
hopx
zgc:123068
igsf9a
krt1−c5
bach1b
rac2
si:ch211−103f14.3
retsat
msi2b
sepp1a
sox1a
si:ch211−266a5.12
zdhhc13
si:dkey−17m8.1
bpnt1
gstm.1
itpk1b
rab3aa
mipol1
hdac6
mhc1zba
slc25a22
copz2
anxa4
fermt1
aldh2.1
FO681288.1
rhbdf1a
snx14
cyp2v1
cox6b2
net1
rab5b
cdh1
fam45a
derl2
si:ch211−127l15.5
ctnnal1
ptprga
dvl2
elovl1a
slc35b1
rbfox2
grp
kat6a
GOLM1
gpx1b
zdhhc16b
spint2
ATP5J.1
rnf185
mical2b
smad5
edem2
rab5ab
mcf2lb
gigyf1
farp1
eef1da
cav2
rgs12a
eya2
bambia
wu:fb18f06
sox21a
tfdp1a
F8A2
col4a6
ndrg3b
stam2
ywhabl
CT956002.1
bc2
csrnp1a
selm
mrpl4
adam10a
gbgt1l3
fndc7rs4
tspan1
crip1
s100s
tuba8l
VPS37D
pmp22a
emc10
hmp19
six2a
frya
si:dkey−182g1.2
birc2
Expression
2
1
0
−1
−2

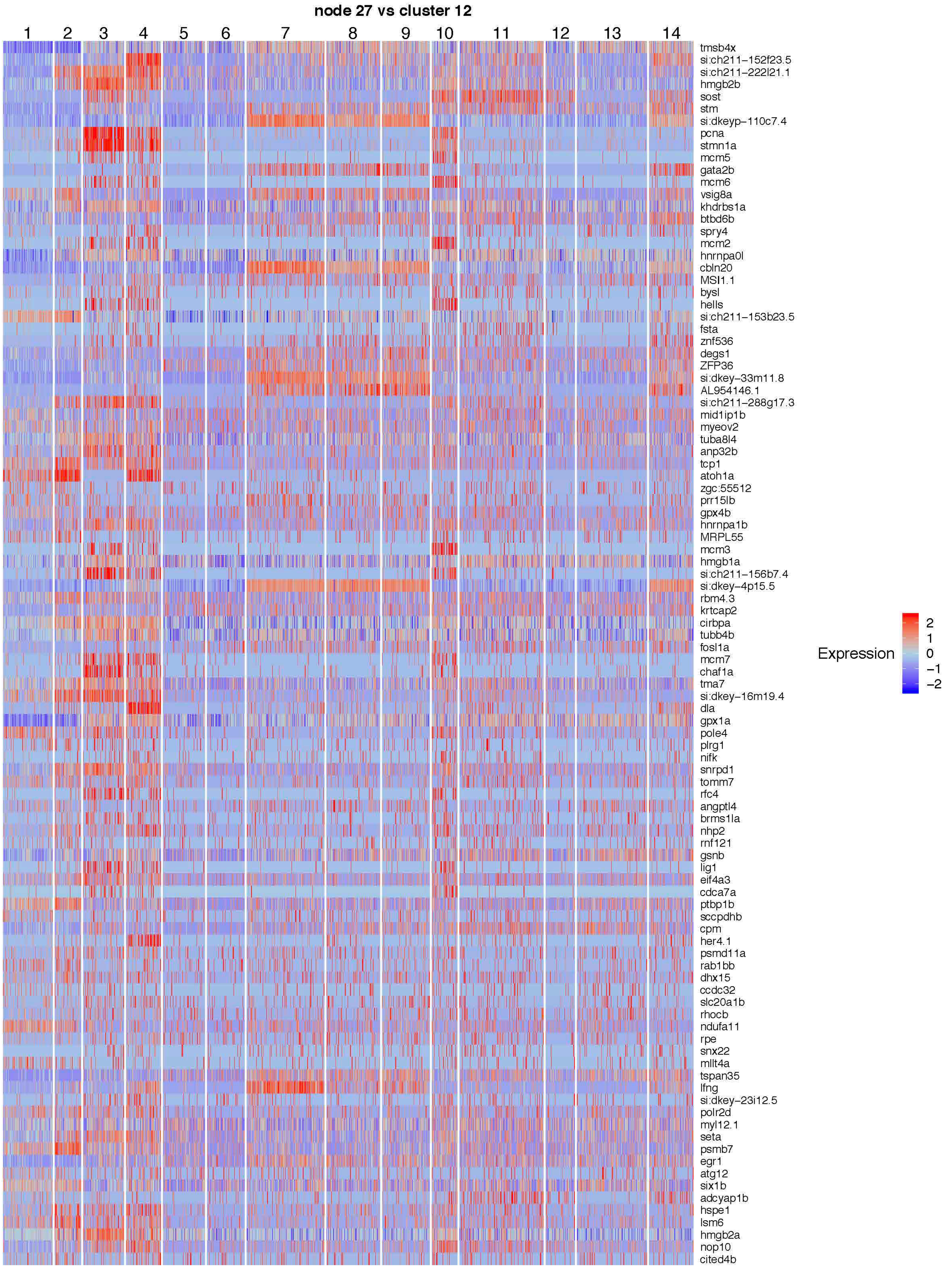
node 27 vs cluster 12
1
2
3
4
5
6
7
8
9
10
11
12
13
14
tmsb4x
si:ch211-152f23.5
si:ch211-222l21.1
hmgb2b
sost
stm
si:dkeyp-110c7.4
pcna
stmn1a
mcm5
gata2b
mcm6
vsig8a
khdrbs1a
btbd6b
spry4
mcm2
hnrnpa0l
cbln20
MSI1.1
bysl
hells
si:ch211-153b23.5
fsta
znf536
degs1
ZFP36
si:dkey-33m11.8
AL954146.1
si:ch211-288g17.3
mid1ip1b
myeov2
tuba8l4
anp32b
tcp1
atoh1a
zgc:55512
prr15lb
gpx4b
hnrnpa1b
MRPL55
mcm3
hmgb1a
si:ch211-156b7.4
si:dkey-4p15.5
rbm4.3
krtcap2
cirbpa
tubb4b
fosl1a
mcm7
chaf1a
tma7
si:dkey-16m19.4
dla
gpx1a
pole4
plrg1
nifk
snrpd1
tomm7
rfc4
angptl4
brms1la
nhp2
rnf121
gsnb
lig1
eif4a3
cdca7a
ptbp1b
sccpdhb
cpm
her4.1
psmd11a
rab1bb
dhx15
ccdc32
slc20a1b
rhocb
ndufa11
rpe
snx22
mllt4a
tspan35
lfng
si:dkey-23i12.5
polr2d
myl12.1
seta
psmb7
egr1
atg12
six1b
adcyap1b
hspe1
lsm6
hmgb2a
nop10
cited4b
Expression
2
1
0
−1
−2

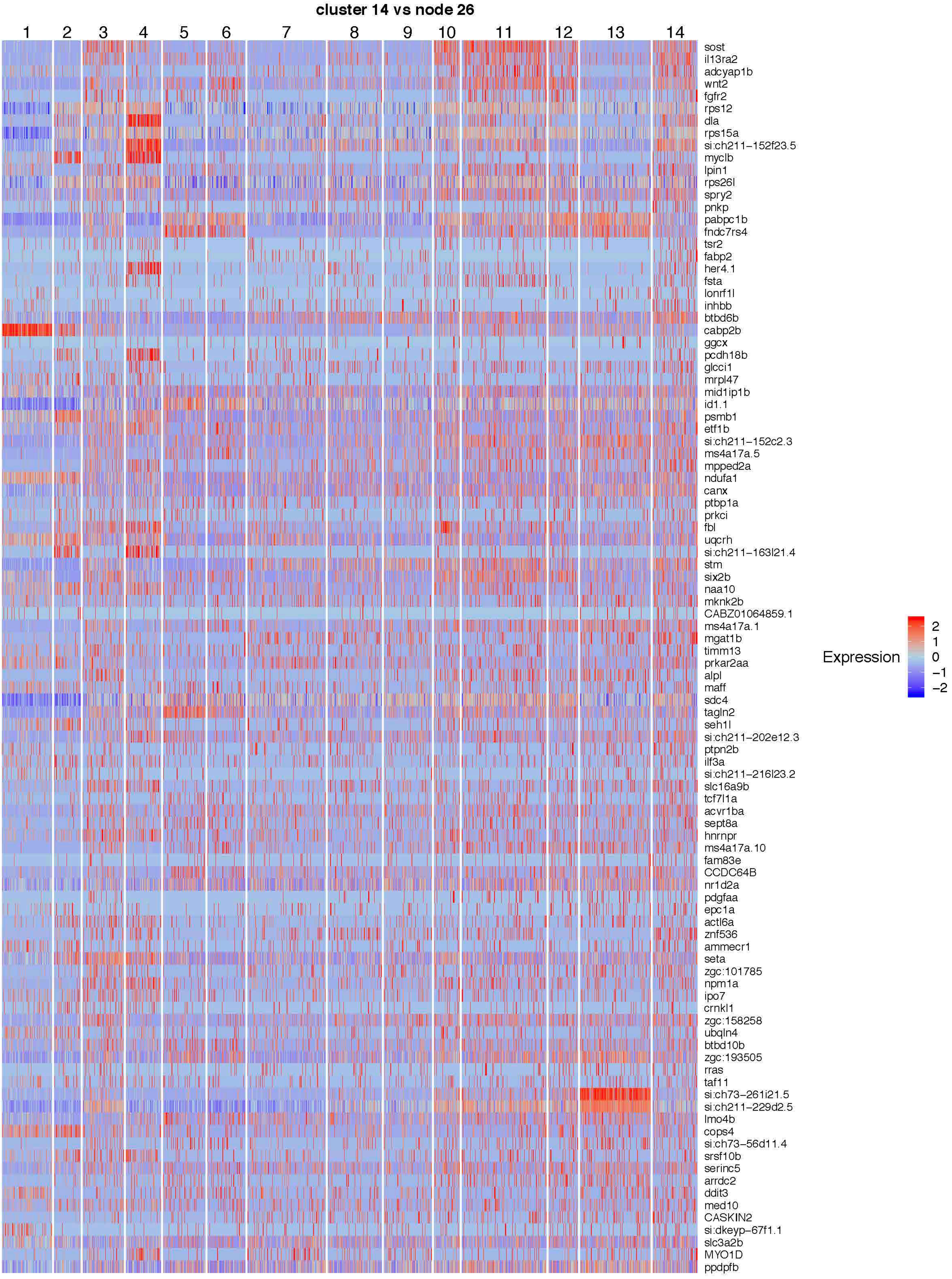
cluster 14 vs node 26
1
2
3
4
5
6
7
8
9
10
11
12
13
14
sost
il13ra2
adcyap1b
wnt2
fgfr2
rps12
dla
rps15a
si:ch211−152f23.5
myclb
lpin1
rps26l
spry2
pnkp
pabpc1b
fndc7rs4
tsr2
fabp2
her4.1
fsta
lonrf1l
inhbb
btbd6b
cabp2b
ggcx
pcdh18b
glcci1
mrpl47
mid1ip1b
id1.1
psmb1
etf1b
si:ch211−152c2.3
ms4a17a.5
mpped2a
ndufa1
canx
ptbp1a
prkci
fbl
uqcrh
si:ch211−163l21.4
stm
six2b
naa10
mknk2b
CABZ01064859.1
ms4a17a.1
mgat1b
timm13
prkar2aa
alpl
maff
sdc4
tagln2
seh1l
si:ch211−202e12.3
ptpn2b
ilf3a
si:ch211−216l23.2
slc16a9b
tcf7l1a
acvr1ba
sept8a
hnrnpr
ms4a17a.10
fam83e
CCDC64B
nr1d2a
pdgfaa
epc1a
actl6a
znf536
ammecr1
seta
zgc:101785
npm1a
ipo7
crnkl1
zgc:158258
ubqln4
btbd10b
zgc:193505
rras
taf11
si:ch73−261i21.5
si:ch211−229d2.5
lmo4b
cops4
si:ch73−56d11.4
srsf10b
serinc5
arrdc2
ddit3
med10
CASKIN2
si:dkeyp−67f1.1
slc3a2b
MYO1D
ppdpfb
Expression
2
1
0
−1
−2

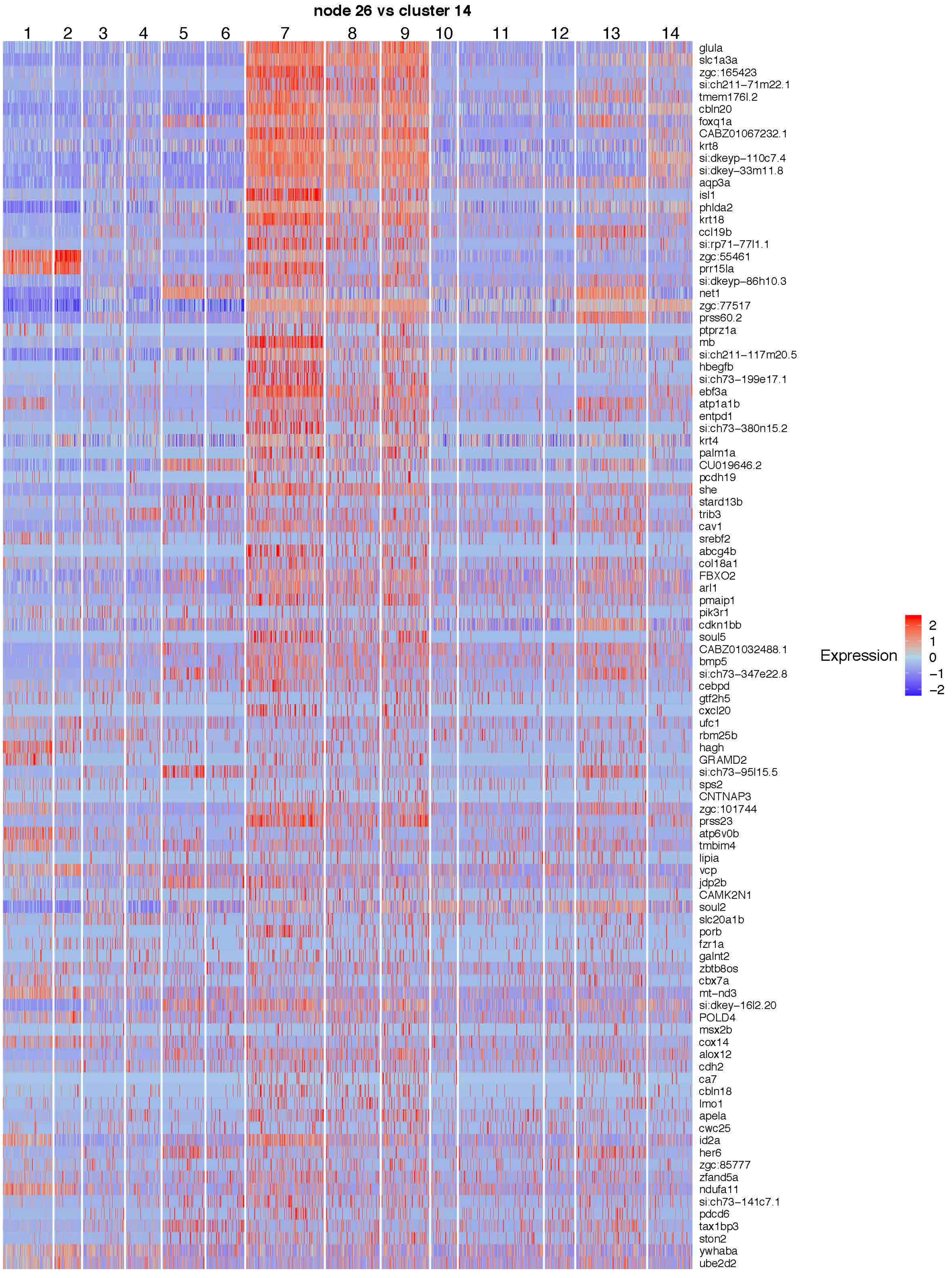
node 26 vs cluster 14
1
2
3
4
5
6
7
8
9
10
11
12
13
14
glula
slc1a3a
zgc:165423
si:ch211−71m22.1
tmem176l.2
cbln20
foxq1a
CABZ01067232.1
krt8
si:dkeyp−110c7.4
si:dkey−33m11.8
aqp3a
isl1
phlda2
krt18
ccl19b
si:rp71−77l1.1
zgc:55461
prr15la
si:dkeyp−86h10.3
net1
zgc:77517
prss60.2
ptprz1a
mb
si:ch211−117m20.5
hbegfb
si:ch73−199e17.1
ebf3a
atp1a1b
entpd1
si:ch73−380n15.2
krt4
palm1a
CU019646.2
pcdh19
she
stard13b
trib3
cav1
srebf2
abcg4b
col18a1
FBXO2
arl1
pmaip1
pik3r1
cdkn1bb
soul5
CABZ01032488.1
bmp5
si:ch73−347e22.8
cebpd
gtf2h5
cxcl20
ufc1
rbm25b
hagh
GRAMD2
si:ch73−95l15.5
sps2
CNTNAP3
zgc:101744
prss23
atp6v0b
tmbim4
lipia
vcp
jdp2b
CAMK2N1
soul2
slc20a1b
porb
fzr1a
galnt2
zbtb8os
cbx7a
mt−nd3
si:dkey−16l2.20
POLD4
msx2b
cox14
alox12
cdh2
ca7
cbln18
lmo1
apela
cwc25
id2a
her6
zgc:85777
zfand5a
ndufa11
si:ch73−141c7.1
pdcd6
tax1bp3
ston2
ywhaba
ube2d2
Expression
2
1
0
−1
−2

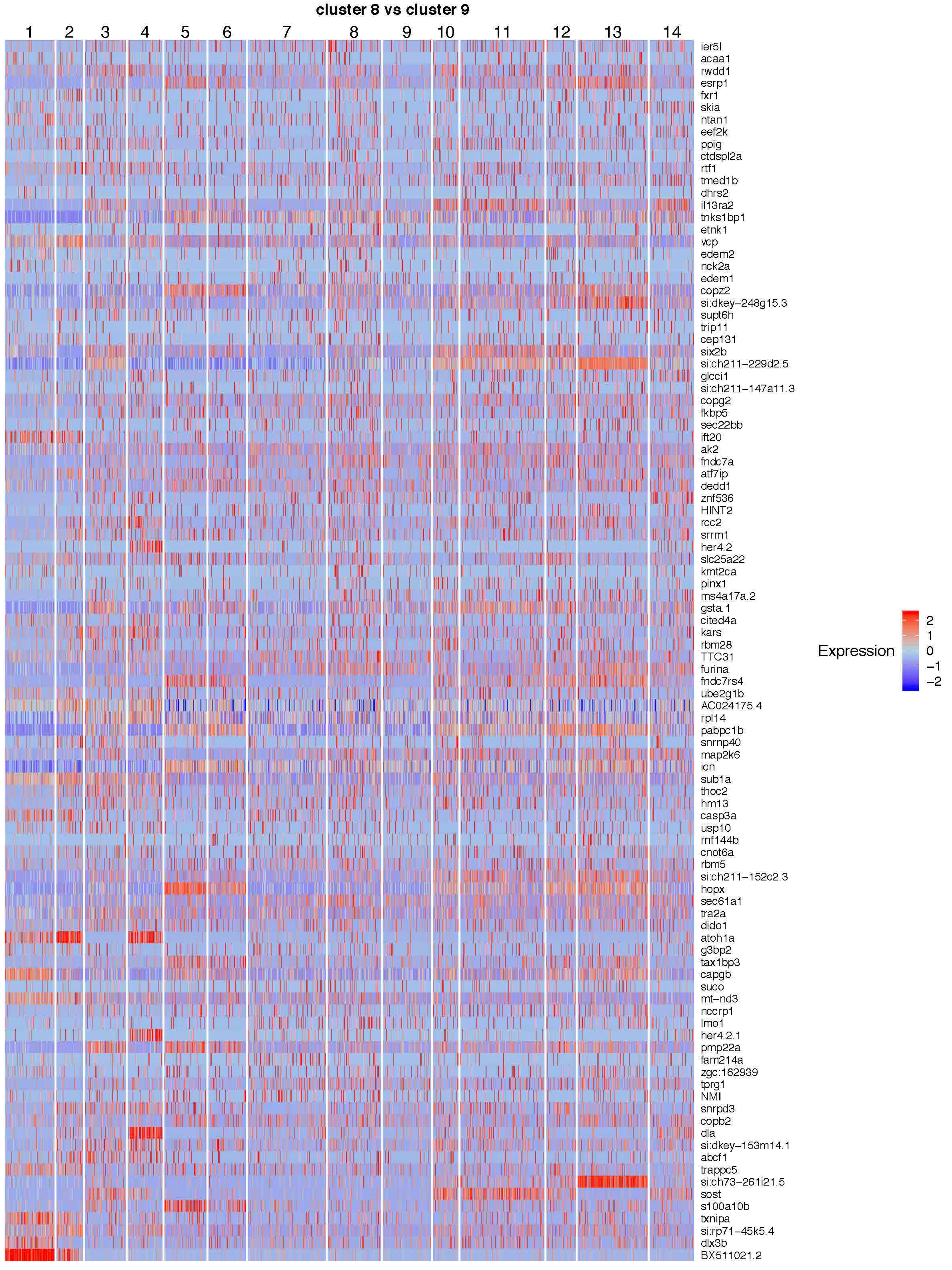

cluster 8 vs cluster 9
1
2
3
4
5
6
7
8
9
10
11
12
13
14
ier5l
acaa1
rwdd1
esrp1
fxr1
skia
ntan1
eef2k
ppig
ctdspl2a
rtf1
tmed1b
dhrs2
il13ra2
tnks1bp1
etnk1
vcp
edem2
nck2a
edem1
copz2
si:dkey−248g15.3
supt6h
trip11
cep131
six2b
si:ch211−229d2.5
glcci1
si:ch211−147a11.3
copg2
fkbp5
sec22bb
ift20
ak2
fndc7a
atf7ip
dedd1
znf536
HINT2
rcc2
srrm1
her4.2
slc25a22
kmt2ca
pinx1
ms4a17a.2
gsta.1
cited4a
kars
rbm28
TTC31
furina
fndc7rs4
ube2g1b
AC024175.4
rpl14
pabpc1b
snrnp40
map2k6
icn
sub1a
thoc2
hm13
casp3a
usp10
rnf144b
cnot6a
rbm5
si:ch211−152c2.3
hopx
sec61a1
tra2a
dido1
atoh1a
g3bp2
tax1bp3
capgb
suco
mt−nd3
nccrp1
lmo1
her4.2.1
pmp22a
fam214a
zgc:162939
tprg1
NMI
snrpd3
copb2
dla
si:dkey−153m14.1
abcf1
trappc5
si:ch73−261i21.5
sost
s100a10b
txnipa
si:rp71−45k5.4
dlx3b
BX511021.2
Expression
2
1
0
−1
−2

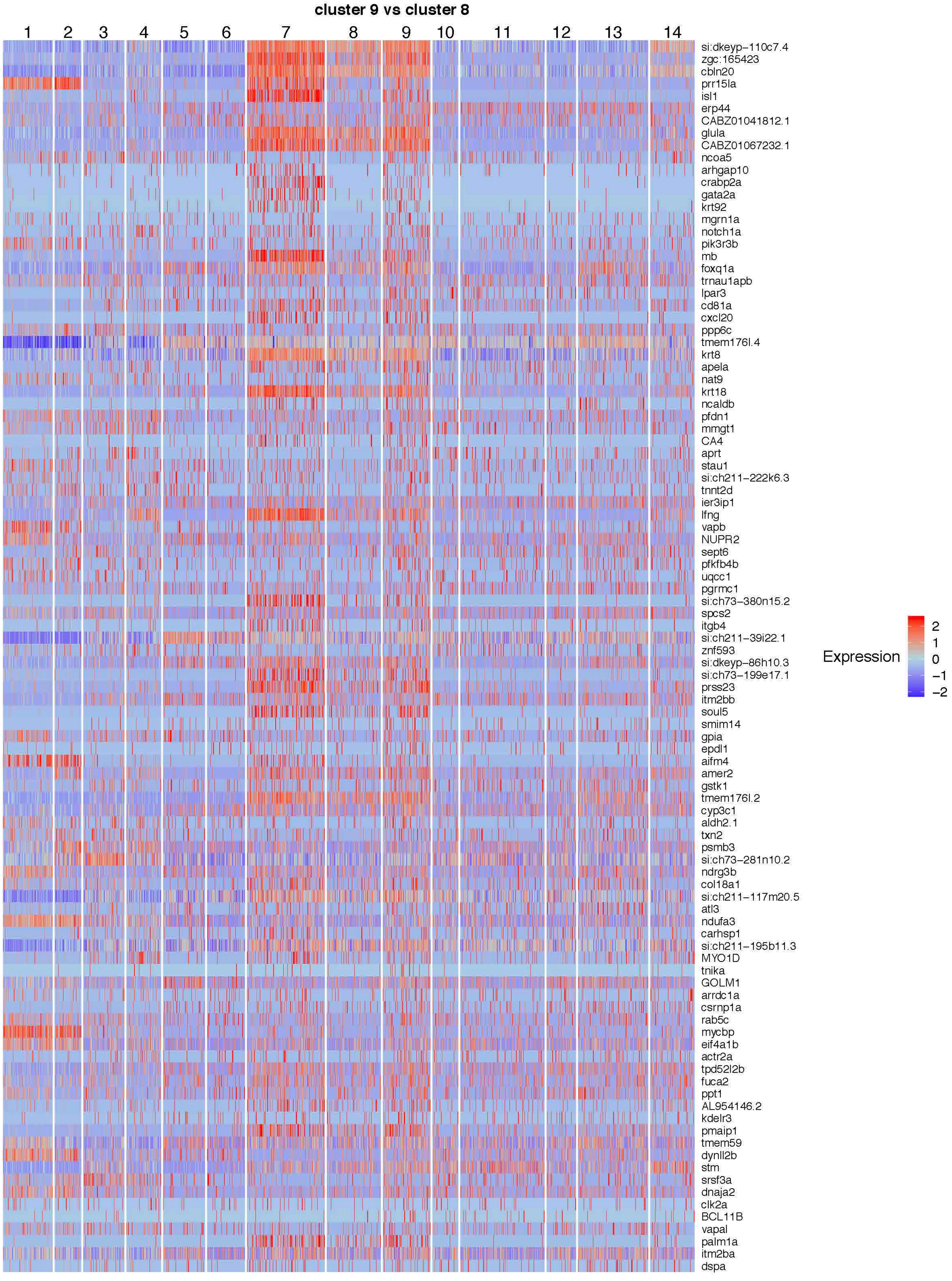
cluster 9 vs cluster 8
1
2
3
4
5
6
7
8
9
10
11
12
13
14
si:dkeyp−110c7.4
zgc:165423
cbln20
prr15la
isl1
erp44
CABZ01041812.1
glula
CABZ01067232.1
ncoa5
arhgap10
crabp2a
gata2a
krt92
mgrn1a
notch1a
pik3r3b
mb
foxq1a
trnau1apb
lpar3
cd81a
cxcl20
ppp6c
tmem176l.4
krt8
apela
nat9
krt18
ncaldb
pfdn1
mmgt1
CA4
aprt
stau1
si:ch211−222k6.3
tnnt2d
ier3ip1
lfng
vapb
NUPR2
sept6
pfkfb4b
uqcc1
pgrmc1
si:ch73−380n15.2
spcs2
itgb4
si:ch211−39i22.1
znf593
si:dkeyp−86h10.3
si:ch73−199e17.1
prss23
itm2bb
soul5
smim14
gpia
epdl1
aifm4
amer2
gstk1
tmem176l.2
cyp3c1
aldh2.1
txn2
psmb3
si:ch73−281n10.2
ndrg3b
col18a1
si:ch211−117m20.5
atl3
ndufa3
carhsp1
si:ch211−195b11.3
MYO1D
tnika
GOLM1
arrdc1a
csrnp1a
rab5c
mycbp
eif4a1b
actr2a
tpd52l2b
fuca2
ppt1
AL954146.2
kdelr3
pmaip1
tmem59
dynll2b
stm
srsf3a
dnaja2
clk2a
BCL11B
vapal
palm1a
itm2ba
dspa
Expression
2
1
0
−1
−2

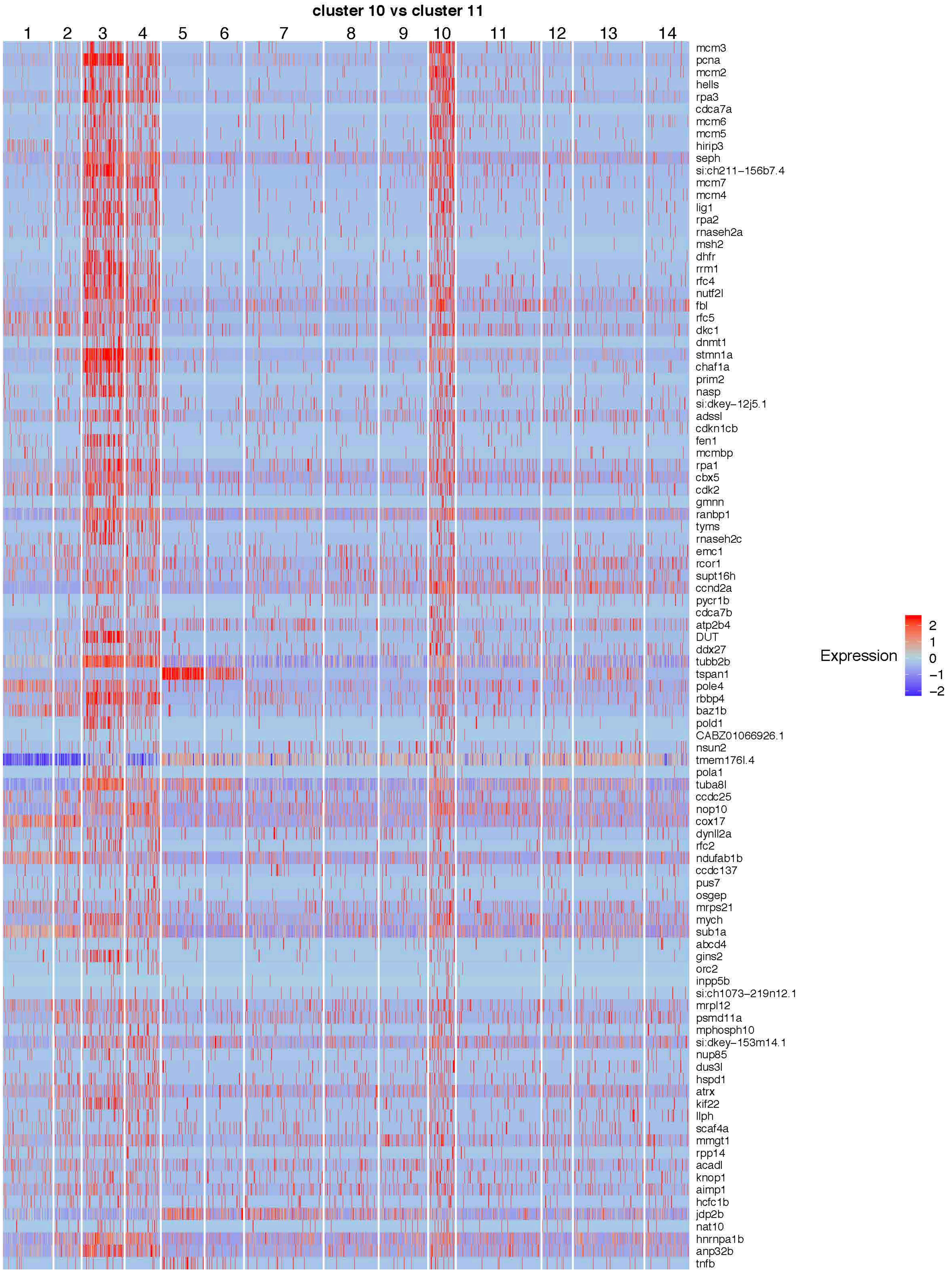
cluster 10 vs cluster 11
1
2
3
4
5
6
7
8
9
10
11
12
13
14
mcm3
pcna
mcm2
hells
rpa3
cdca7a
mcm6
mcm5
hirip3
seph
si:ch211−156b7.4
mcm7
mcm4
lig1
rpa2
rnaseh2a
msh2
dhfr
rrm1
rfc4
nutf2l
fbl
rfc5
dkc1
dnmt1
stmn1a
chaf1a
prim2
nasp
si:dkey−12j5.1
adssl
cdkn1cb
fen1
mcmbp
rpa1
cbx5
cdk2
gmnn
ranbp1
tyms
rnaseh2c
emc1
rcor1
supt16h
ccnd2a
pycr1b
cdca7b
atp2b4
DUT
ddx27
tubb2b
tspan1
pole4
rbbp4
baz1b
pold1
CABZ01066926.1
nsun2
tmem176l.4
pola1
tuba8l
ccdc25
nop10
cox17
dynll2a
rfc2
ndufab1b
ccdc137
pus7
osgep
mrps21
mych
sub1a
abcd4
gins2
orc2
inpp5b
si:ch1073−219n12.1
mrpl12
psmd11a
mphosph10
si:dkey−153m14.1
nup85
dus3l
hspd1
atrx
kif22
llph
scaf4a
mmgt1
rpp14
acadl
knop1
aimp1
hcfc1b
jdp2b
nat10
hnrnpa1b
anp32b
tnfb
Expression
2
1
0
−1
−2

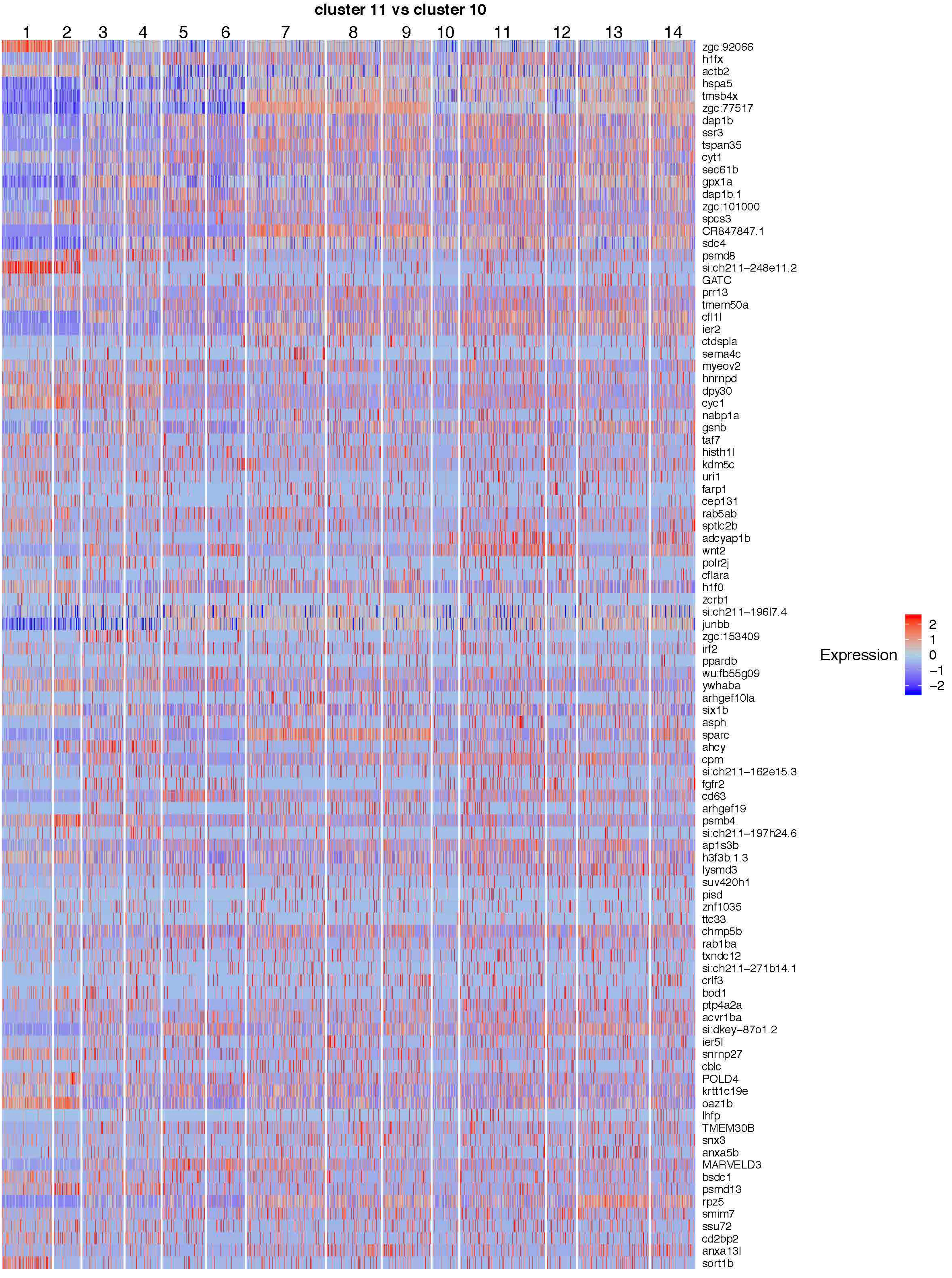
cluster 11 vs cluster 10
1
2
3
4
5
6
7
8
9
10
11
12
13
14
zgc:92066
h1fx
actb2
hspa5
tmsb4x
zgc:77517
dap1b
ssr3
tspan35
cyt1
sec61b
gpx1a
dap1b.1
zgc:101000
spcs3
CR847847.1
sdc4
psmd8
si:ch211−248e11.2
GATC
prr13
tmem50a
cfl1l
ier2
ctdspla
sema4c
myeov2
hnrnpd
dpy30
cyc1
nabp1a
gsnb
taf7
histh1l
kdm5c
uri1
farp1
cep131
rab5ab
sptlc2b
adcyap1b
wnt2
polr2j
cflara
h1f0
zcrb1
si:ch211−196l7.4
junbb
zgc:153409
irf2
ppardb
wu:fb55g09
ywhaba
arhgef10la
six1b
asph
sparc
ahcy
cpm
si:ch211−162e15.3
fgfr2
cd63
arhgef19
psmb4
si:ch211−197h24.6
ap1s3b
h3f3b.1.3
lysmd3
suv420h1
pisd
znf1035
ttc33
chmp5b
rab1ba
txndc12
si:ch211−271b14.1
crlf3
bod1
ptp4a2a
acvr1ba
si:dkey−87o1.2
ier5l
snrnp27
cblc
POLD4
krtt1c19e
oaz1b
lhfp
TMEM30B
snx3
anxa5b
MARVELD3
bsdc1
psmd13
rpz5
smim7
ssu72
cd2bp2
anxa13l
sort1b
Expression
2
1
0
−1
−2
